# Optimal carbon partitioning reconciles the apparent divergence between optimal and observed canopy profiles of photosynthetic capacity

**DOI:** 10.1101/2020.09.16.300202

**Authors:** Thomas N. Buckley

## Abstract

**Research conducted:** Photosynthetic capacity per unit irradiance is greater, and the marginal carbon revenue of water (∂*A*/∂*E*) is smaller, in shaded leaves than sunlit leaves, apparently contradicting optimization theory. I tested the hypothesis that these patterns arise from optimal carbon partitioning subject to biophysical constraints on leaf water potential.

**Methods:** In a whole plant model with two canopy modules, I adjusted carbon partitioning, nitrogen partitioning and leaf water potential to maximize carbon profit or canopy photosynthesis, and recorded how gas exchange parameters compared between shaded and sunlit modules in the optimum.

**Key results:** The model predicted that photosynthetic capacity per unit irradiance should be larger, and ∂*A*/∂*E* smaller, in shaded modules compared to sunlit modules. This was attributable partly to radiation-driven differences in evaporative demand, and partly to differences in hydraulic conductance arising from the need to balance marginal returns on stem carbon investment between modules. The model verified, however, that invariance in the marginal carbon revenue of N (∂*A*/∂*N*) is in fact optimal.

**Conclusion:** The Cowan-Farquhar optimality solution (invariance of ∂*A*/∂*E*) does not apply to spatial variation within a canopy. The resulting variation in carbon-water economy explains differences in capacity per unit irradiance, reconciling optimization theory with observations.

## Introduction

Scaling photosynthesis and transpiration from leaves to canopies is made difficult by wide spatial variation among canopy locations in key parameters that determine gas exchange, particularly photosynthetic capacity and stomatal conductance. Modelers commonly use optimization theory – the hypothesis that plants have evolved to maximize the return on investment of limiting resources – to infer canopy profiles of gas exchange parameters (Amthor, 1994; de Pury & Farquhar, 1997). Applied to canopy photosynthesis, optimization theory predicts that a fixed total supply of photosynthetic nitrogen is optimally distributed when the marginal carbon product of N, ∂*A*/∂*N*, is invariant among canopy positions and among functional N pools within each location (Field, 1983):

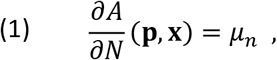

where *A* is net CO_2_ assimilation rate, averaged over some period, such as one day, during which it is assumed that N cannot be redistributed among pools or locations; **p** and **x** denote vectors of functional N pools (carboxylation, regeneration and light harvesting) and canopy positions, respectively; and *μ*_n_ is a Lagrange multiplier that is invariant among N pools and canopy positions (a list of symbols is given in Table 1). When Eqn 1 is applied to simple models of canopy gas exchange, it predicts that photosynthetic capacity should vary among canopy locations in proportion to the average daily or seasonal irradiance (Field, 1983; Hirose & Werger, 1987; Farquhar, 1989; Sands, 1995), or equivalently, that the ratio of photosynthetic capacity between any two canopy layers should be equal to the ratio of irradiance between the layers. That prediction is very useful for upscaling models of leaf photosynthesis, because it allows an unknown biological property (the spatial distribution of photosynthetic capacity in a canopy) to be inferred from a more easily measured and/or simulated physical property (the distribution of light). Under certain limiting conditions, it even makes leaf-scale models of photosynthesis “scale-invariant”, meaning that the model works whether applied using leaf-level parameters or their canopy-level averages (Farquhar, 1989).

**Table 1.**
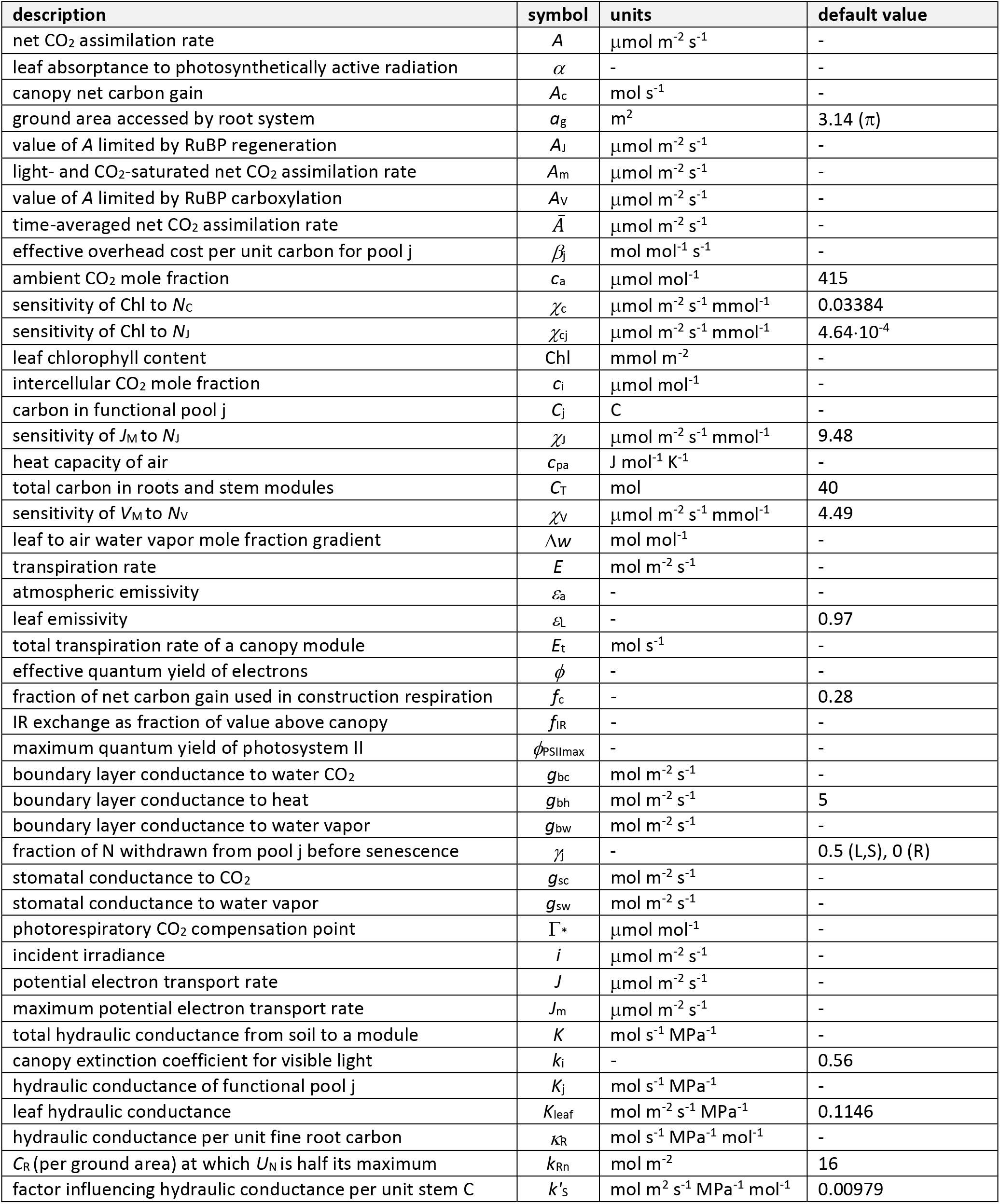
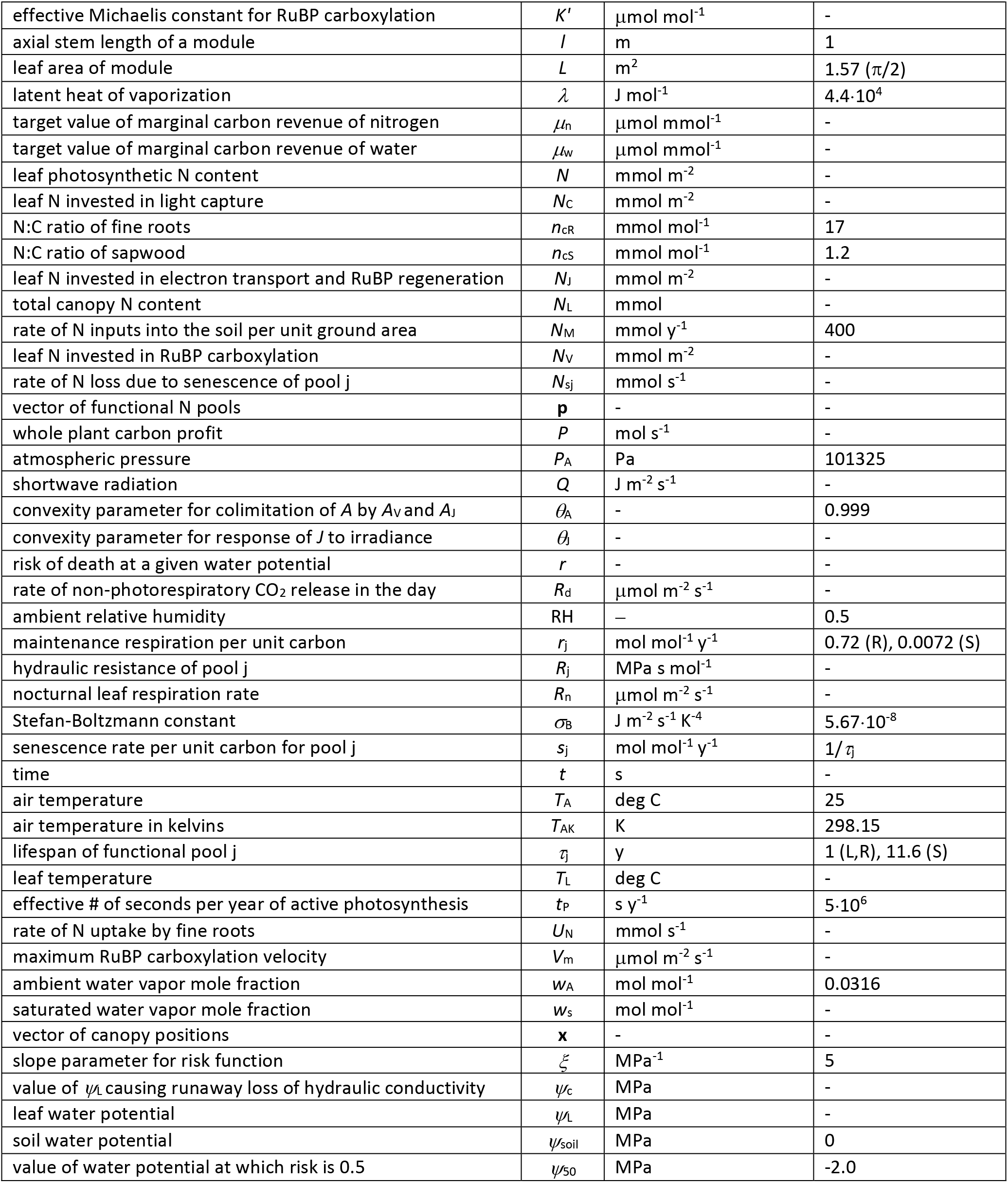
Mathematical symbols, units and default values; annotations in the default value column refer to functional pools (L: leaves; S: stem [sapwood]; R: roots).

Those predictions from optimality theory contrast starkly with observations. Abundant data across many species and functional types show that the ratio of photosynthetic capacity between shaded and sunlit layers systematically exceeds the ratio of irradiance; that is, more sunlit canopy locations have less photosynthetic capacity per unit irradiance (e.g., Hirose & Werger, 1987; Evans, 1993; Hollinger, 1996; de Pury & Farquhar, 1997; Makino *et al.*, 1997; Bond *et al.*, 1999; Friend, 2001; Frak *et al.*, 2002; Kull, 2002; Lloyd *et al.*, 2010; Niinemets *et al.*, 2015; Hikosaka *et al.*, 2016; Salter *et al.*, 2020). No single hypothesis seems adequate to explain this apparent divergence between theory and observations across all environments and functional types (Niinemets, 2012; Buckley *et al.*, 2013; Niinemets *et al.*, 2015; Hikosaka *et al.*, 2016) – posing a puzzle for physiologists and ecologists, and casting doubt on the theory and its utility for predicting and interpreting plant function.

Most theoretical studies of canopy N partitioning have used models in which spatial variation in photosynthesis is driven solely by patterns of N and light, and have thus overlooked the potential influence of spatial patterns of water loss on photosynthetic N economy (Buckley *et al.*, 2002). It has been long known that stomatal conductance is often systematically suppressed in upper-canopy leaves (Ryan & Yoder, 1997; Delzon *et al.*, 2004; Koch *et al.*, 2004). Such suppression would cause optimal photosynthetic capacity to be lower than expected in the upper canopy (Peltoniemi *et al.*, 2012; Buckley *et al.*, 2013), and could arise from low leaf water potentials – caused either by the greater hydraulic resistance encountered in transporting water to more distal sites in the canopy, or from elevated evaporative demand in more sunlit locations (Ambrose *et al.*, 2016; Bachofen *et al.*, 2020). Indeed, stomatal conductance responds negatively both to reduced water potential and increased evaporative demand (Buckley, 2019).

Yet the empirical fact of reduced stomatal conductance in the upper canopy does not by itself resolve the apparent failure of optimization theory. Why would a plant not simply provide sunlit leaves with greater capacity for water transport, to prevent reductions in water potential – and hence stomatal conductance and optimal photosynthetic capacity – resulting from height, transport distance, or evaporative demand? Peltoniemi et al. (2012) found that stomatal conductance and photosynthetic capacity should not in fact be suppressed in the upper canopy if hydraulic conductance is optimally distributed between canopy modules, suggesting that hydraulic conductance is not optimally distributed in real plants. Similarly, Buckley et al. (2014) found that the spatial distribution of stomatal conductance and water loss was systematically suboptimal in grapevine canopies, with sunlit leaves transpiring less than predicted and shaded leaves transpiring more.

The latter result was premised on the same logic as Equation 1: namely, if one assumes that a given total amount of water loss is available for distribution in the canopy, then, provided water use earns carbon gain with diminishing returns (∂^2^*A*/∂*E*^2^ < 0), canopy carbon gain is maximized if the marginal carbon revenue of water ((∂*A*/∂*g*_sw_)/(∂*E*/∂*g*_sw_) ≡ ∂*A*/∂*E*) is invariant and equal to a Lagrange multiplier, *μ*_w_:

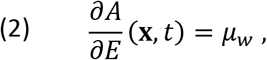

where *E* is transpiration rate, *g*_sw_ is stomatal conductance to H_2_O, and *t* is time (Buckley *et al.*, 2002). (Cowan and Farquhar (1977) derived a result identical to Eqn 2; although they focused on invariance of ∂*A*/∂*E* over time, rather than among leaves in a canopy, the domain of variation does not affect the mathematical validity of the solution; Buckley et al. 2002). The conclusion of Peltoniemi et al. (2012) was likewise premised on the assumption that the total amount of hydraulic conductance is an imposed constraint, such that ∂*A*/∂*K* should be invariant in the optimum.

My objective here was to determine whether those assumptions are consistent with a broader interpretation of optimization theory, in which the problem is extended to a higher level of organization: namely, optimal carbon partitioning at the whole-plant level, subject to biophysical and economic constraints on leaf water potential. I used simulations from a whole-plant model to test the hypothesis that spatial invariance in ∂*A*/∂*N*, ∂*A*/∂*E* and ∂*A*/∂*K*, and by extension invariance in optimal photosynthetic capacity per unit incident irradiance between sunlit and shaded regions of the canopy, does in fact emerge from optimal carbon partitioning.

## Description

I simulated canopy photosynthesis in an imaginary plant consisting of two canopy modules (one “sunlit” and one “shaded”, the latter with lower leaf-level incident irradiance than the former) and a root system. Each canopy module a stem carbon pool and a fixed amount of leaf area. I assigned each module three other parameters: incident irradiance (*i*), boundary layer conductance to heat (*g*_bh_) and axial stem length (*l*); air temperature and relative humidity were assumed to be identical between the two canopy modules. In this model, stem carbon determines each module’s total xylem conducting area and thus stem hydraulic conductance, and root carbon determines the root hydraulic conductance shared by both modules and the total supply of nitrogen available. I maximized either canopy photosynthesis (the sum of photosynthesis in both modules) or carbon profit (canopy photosynthesis minus the amortized carbon costs of maintenance and turnover of the carbon pools) by numerically adjusting (i) the partitioning of a fixed total carbon supply (among the root and stem carbon pools), (ii) the partitioning of available N (between the two canopy modules, and among N pools for RuBP carboxylation, electron transport and light capture in each module), and (iii) the values of leaf water potential in each module. I recorded how gas exchange parameters that emerged from the optimization in each module compared to one another in relation to the ratio of irradiance between the two modules.

### Penalizing the non-stomatal consequences of low leaf water potential

In the carbon balance model summarized above and presented in the Appendix, maintaining a high leaf water potential is never beneficial for carbon gain, because any decrease in *ψ*_L_ leads directly to an increase in stomatal conductance (Eqn A24), and therefore an increase in carbon gain (Eqn A8). The main reason *ψ*_L_ does not generally become arbitrarily low in real plants is that doing so has negative consequences that are independent of stomatal conductance (plants have thus evolved to close stomata at low *ψ*_L_; but such adaptive patterns cannot be taken as prior constraints if the objective, as in this study, is to identify adaptive patterns). Very low *ψ*_L_ leads to irreversible loss of water transport capacity (Tyree & Sperry, 1988, 1989; Choat *et al.*, 2012; McCulloh *et al.*, 2019) with consequent runaway desiccation. Although some non-stomatal consequences of low *ψ*_L_, such as depression of photosynthetic capacity, can manifest directly in reduced photosynthesis (Lawlor & Tezara, 2009), the weight of empirical evidence suggests such effects are generally not substantial until water potential is already low enough to cause both stomatal closure and cavitation (Kaiser, 1987; Downton *et al.*, 1988; Sharkey & Seemann, 1989; Quick *et al.*, 1992; Centritto *et al.*, 2003; Koch *et al.*, 2004; Chaves *et al.*, 2009). As a result, most non-stomatal effects of low *ψ*_L_ on carbon balance are intrinsically probabilistic – they are driven by the risk of exceeding the threshold for runaway cavitation (*ψ*_c_), rather than by immediate short-term carbon costs – so they influence the expected value of total carbon gain over the module’s lifespan, 〈*A*〉, rather than the instantaneous assimilation rate, *A*. For these reasons, I modeled non-stomatal costs of low *ψ*_L_ as a non-dimensional *risk factor* that is a function of *ψ*_L_ and is multiplied by the photosynthesis rate calculated in the absence of non-stomatal effects of *ψ*_L_:

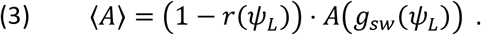

I assumed the risks represented by *r* varied sigmoidally with *ψ*_L_, increasing from zero in an accelerating as *ψ*_L_ declines from zero towards a threshold value, *ψ*_50_, at which *r* = 0.5, and then then decelerating as *ψ*_L_ declines further. A convenient function with these properties is

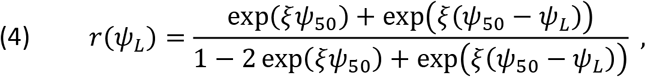

where *ξ* controls the slope of the function at *ψ*_L_ = *ψ*_50_ (large *ξ* = steep slope). The formulation represented by Eqns 3–4 is very similar to those adopted in recent optimization based models of stomatal conductance (e.g., Wolf *et al.*, 2016; Sperry *et al.*, 2017; Eller *et al.*, 2020). However, Eqn 4 is not the hydraulic vulnerability curve; instead, it represents the risk of catastrophic desiccation posed by allowing *ψ*_L_ to reach a given value. That risk is influenced not only by the vulnerability curve but also by factors that affect the likelihood of transiently exceeding *ψ*_c_, such as how quickly stomata can respond to fluctuations in evaporative demand and the probability distribution of such fluctuations.

### Simulations

The model contains nine degrees of freedom, represented by two carbon partitioning fractions, five nitrogen partitioning fractions, and one value of leaf water potential for each canopy module. I adjusted these nine parameters numerically using Solver in Microsoft Excel to maximize either whole-plant carbon profit or total photosynthesis (in each case using the expected values of assimilation rate given by Eqn 3). I repeated this procedure for a range of parameter combinations (Table 2). These included three scenarios for the steepness of the risk curve (controlled by the parameter *ξ* in Eqn 4): two finite values of *ξ* (1 and 5 MPa^−1^) and one scenario representing the limit of large *ξ*. In the latter scenario, 〈*A*〉 was by definition greatest in the limit of *ψ*_L_ → *ψ*_50_ (because r was constant and equal to 1 for *ψ*_L_ ≥ *ψ*_50_, and equal to zero for *ψ*_L_ < *ψ*_50_), so numerical optimization of *ψ*_L_ was unnecessary, and I simply set *ψ*_L_ to *ψ*_50_ and excluded *ψ*_L_ from the list of parameters to optimize. In each simulation, I calculated ∂*A*/∂*E*, ∂*A*/∂*N*, ∂*A*/∂*K* and *∂P/∂N* numerically for each module. I also calculated the light- and CO_2_-saturated assimilation rate (*A*_m_) for each module as the limit of *A* under infinite irradiance and intercellular CO_2_ concentration. The Excel file implementing these calculations is included as Supporting Information Methods S1.

**Table 2.** List of simulations, with values or ranges of parameters adjusted in each.

| simulation | boundary layer conductance ($g_{bh}$) | air temperature ($T_A$) | relative humidity (RH) | hydraulic pathlength of sunlit module ($l_{sunlit}$) | steepness parameter for risk curve ($\xi$) | irradiance ratio ($i_{shaded}/i_{sunlit}$) |
| --- | --- | --- | --- | --- | --- | --- |
| default | **2 mol m<sup>-2</sup> s<sup>-1</sup>** | **25°C** | **50%** | **1 m** | **5 MPa<sup>-1</sup>** | **0.13 – 1.0**<br>($i_{sunlit} = 1500$ $\mu\text{mol m}^{-2} \text{s}^{-1}$;<br>$i_{shaded} = 200 - 1500$) |
| low $g_{bh}$ | **1** | – | – | – | – | |
| high $g_{bh}$ | **1000** | – | – | – | – | |
| low $g_{bh}$ in shaded | **1\*** | – | – | – | – | |
| high $T_A$ | – | **35** | – | – | – | |
| low RH | – | – | **25** | – | – |  |
| long pathlength | – | – | – | **1.5, 2.0** | – |  |
| gradual risk | – | – | – | – | **1** |  |
| steep risk | – | – | – | – | ∞ | |

– : Parameter was set to default value given in first row; \* $g_{bh}$ was 1 and 2 mol m<sup>-2</sup> s<sup>-1</sup> in shaded and sunlit modules, respectively.

### Parameter values and parameter sensitivity

Biological and environmental parameters and their default values are listed in Tables 1 and 2. I estimated biological parameters from a variety of literature sources, as described in Supporting Information Notes S1. I chose values for total carbon supply, soil N input rate, and ground area per module to produce reasonable values for key gas exchange parameters. In the simulations described below, I used the “nominal” values given for each parameter unless otherwise stated.

## Results

When carbon partitioning was optimized in a plant model with two canopy modules (one “sunlit” module with greater irradiance, *i*, than the other “shaded” module), the ratio of photosynthetic capacity between the shaded and sunlit modules exceeded the ratio of irradiances (Fig 1; Table 3 gives values for a number of gas exchange-related parameters for a single simulation in which irradiance in the sunlit module was twice that in the shaded module). In other words, a plot of the capacity ratio on the vertical axis and the irradiance ratio on the horizontal axis systematically diverged above the 1:1 line (thick grey line in Fig 1). This divergence was generally greatest at intermediate and high irradiance ratios. These results held qualitatively regardless of how photosynthetic capacity was quantified (as *V*_m_ or *J*_m_; Fig 1), though the divergence was slightly smaller for *V*_m_ than for *J*_m_. Subsequent results are presented in terms of light- and CO_2_-saturated *A* (*A*_m_).

**Figure 1.**
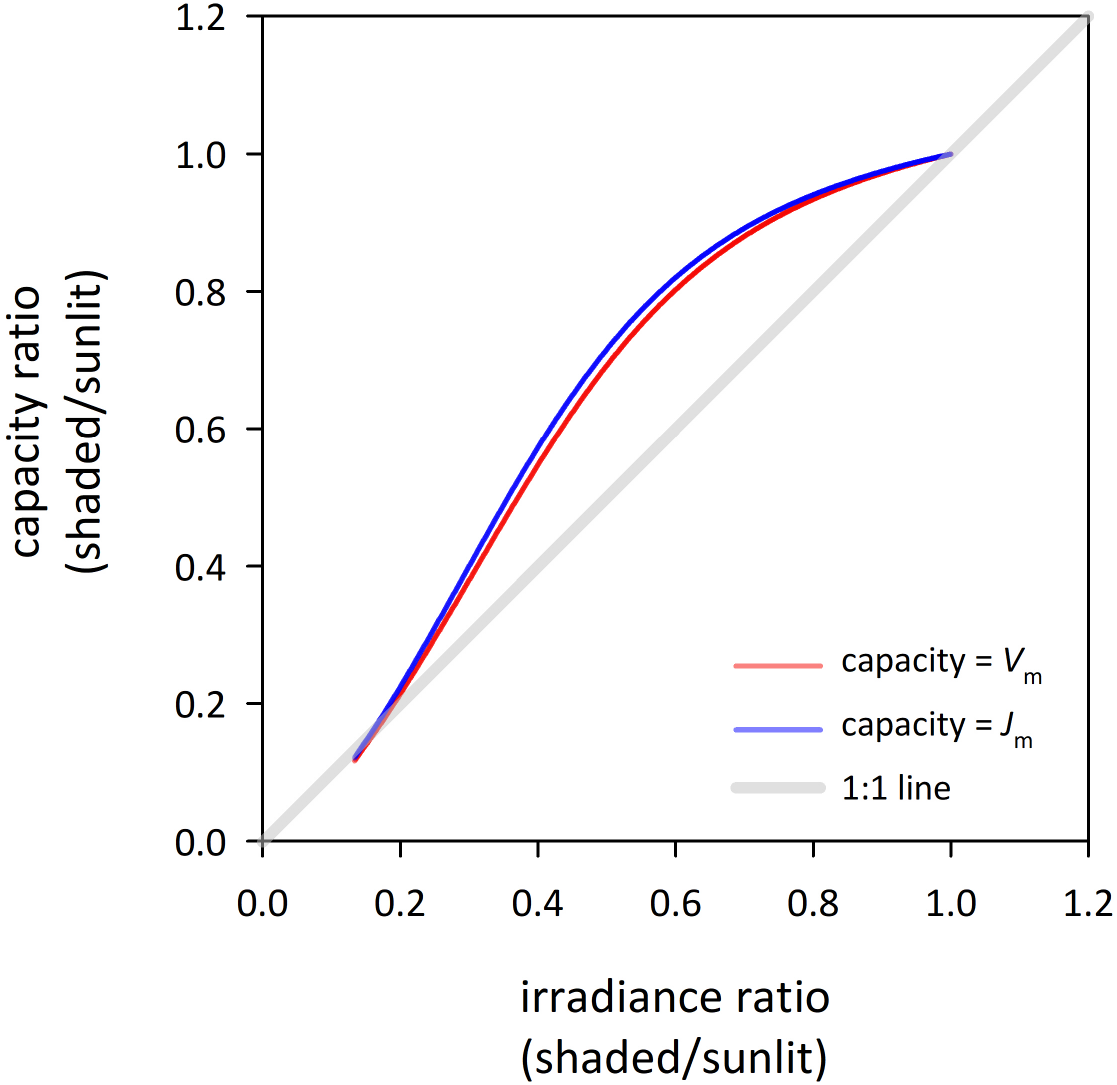
The predicted optimal ratio of photosynthetic capacity (RuBP carboxylation capacity [*V*_m_, red line] or electron transport capacity [*J*_m_, blue line]) between shaded and sunlit modules systematically exceeded the ratio of incident irradiance between the modules, when carbon partitioning was adjusted among roots and stem C pools in both modules so as to maximize whole-plant carbon profit. (The trend for CO_2_- and light-saturated assimilation rate (*A*_m_) was nearly identical to that for *J*_m_, so the two could not be distinguished in a figure and hence only *J*_m_ is not shown. Subsequent figures show results for *A*_m_.)

**Table 3.** Detailed results for example simulation with irradiance ratio = 0.5 (*i* = 1500 and 750 *μ*mol m^−2^ s^−1^ in sunlit and shaded modules, respectively) and default parameters (Table 2); units for ∂*A*/∂*E*, ∂*A*/∂*N*, ∂*P*/∂*N* are μmol mmol^−1^; see Table 1 for other symbols and units.

| variable | sunlit | shaded |
| --- | --- | --- |
| $V_m$ | 119.4 | 82.8 |
| $J_m$ | 188.9 | 135.3 |
| $A_m$ | 46.1 | 33.1 |
| $A$ | 25.2 | 18.1 |
| $c_i$ | 268.8 | 276.3 |
| $g_{sw}$ | 0.310 | 0.227 |
| $E$ | 0.00425 | 0.00317 |
| $T_L$ | 25.05 | 24.91 |
| $\Delta w$ | 0.0157 | 0.0154 |
| $\psi_L$ | -1.46 | -1.37 |
| $\partial A/\partial E$ | 2.20 | 1.83 |
| $\partial A/\partial N$ | 0.328 | 0.318 |
| $\partial P/\partial N$ | 0.201 | 0.201 |

Evaporative demand (Δ*w*) was typically greater in the sunlit module than in the shaded module (Figs 2a,b), due to the greater radiation load in the sunlit module, except at low ambient relative humidity (25% vs the default of 50%) and in some cases at very low irradiance ratios (< 0.3). These differences in Δ*w* moderately increased the positive divergence of the capacity ratio from the irradiance ratio (Figs 2c,d). For example, differences in Δ*w* between the modules were greater when boundary layer conductance (*g*_bh_) was greater in the sunlit module (Fig 2a) or air temperature was increased (Fig 2b), and this translated into a greater divergence of the capacity and irradiance ratios (Figs 2c,d). However, a substantial divergence persisted even if differences in Δ*w* were eliminated by setting the boundary layer conductance to a very large value (ensuring that leaf and air temperatures were equal) (red lines in Figs 2a,c).

**Figure 2.**
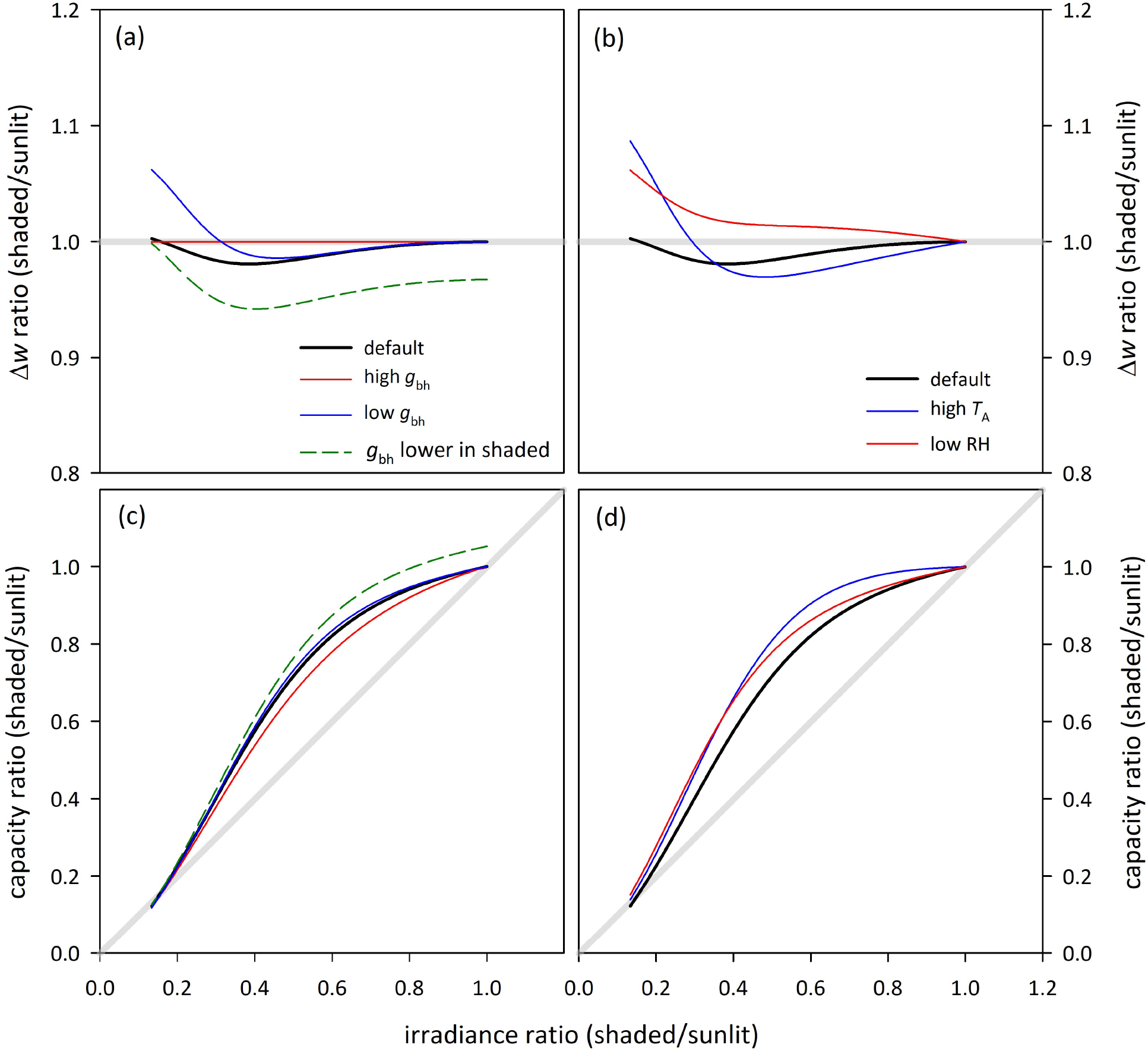
(a,b) The evaporative demand (Δ*w*; leaf to air water vapor mole fraction gradient) was generally lower in the shaded canopy module than in the sunlit module unless leaf temperature was effectively forced to equal air temperature by setting boundary layer conductance to heat to an extremely high value (“high *g*_bh_”, *g*_bh_ = 1000 mol m^−2^ s^−1^). Either reducing *g*_bh_ from 2 (default) to 1 mol m^−2^ s^−1^ (“low *g*_bh_”), setting *g*_bh_ lower in the sunlit module (*g*_bh_ = 1 mol m^−2^ s^−1^, vs. 2 in the sunlit module) or increasing air temperature from *T*_A_ = 25°C (default) to 35°C (“high *T*_A_”) magnified the difference in Δ*w* between the two modules, whereas reducing relative humidity from RH = 50% (default) to 25% (“low RH”) had the opposite effect. (c,d) In most cases, conditions that increased differences in Δ*w* between modules also increased the divergence of the capacity and irradiance ratios. (Capacity ratio is expressed in terms of *A*_m_.) Diagonal grey lines in (c,d) are 1:1 lines.

Simulations in which the sunlit module was hydraulically distal to the shaded module – that is, water was required to travel farther to reach the sunlit module – optimal photosynthetic capacity was predicted to be greater in the shaded module, even if irradiance was as much as 50% greater in the sunlit module (Fig 3).

**Figure 3.**
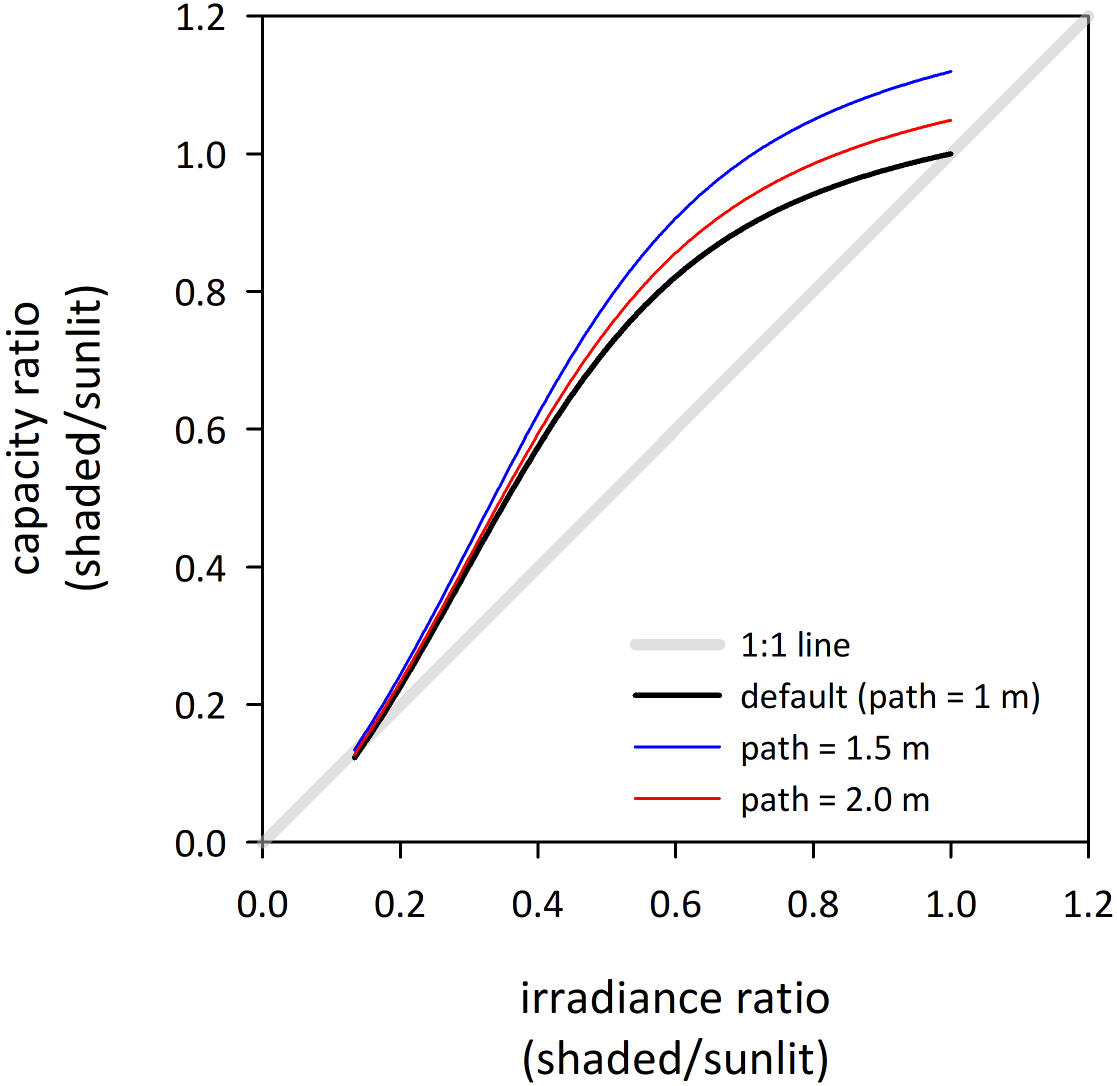
Assigning the sunlit canopy module a greater hydraulic pathlength (*l*_1_ = 1.5 or 2.0 m, blue and red lines, respectively) than the default value (*l*_1_ = 1.0 m, black line) caused the capacity ratio to exceed the irradiance ratio by a greater degree, even if both modules had the same incident irradiance (irradiance ratio = 1.0).

The divergence of the capacity and irradiance ratios was inversely related to the steepness of the risk function used to penalize low leaf water potentials. For example, the divergence was slightly greater if risk was assumed to increase very gradually as *ψ*_L_ declined (Fig 4; inset shows the risk function itself). However, the divergence persisted, except at low irradiance ratios, even if the risk function was infinitely abrupt (Fig 4).

**Figure 4.**
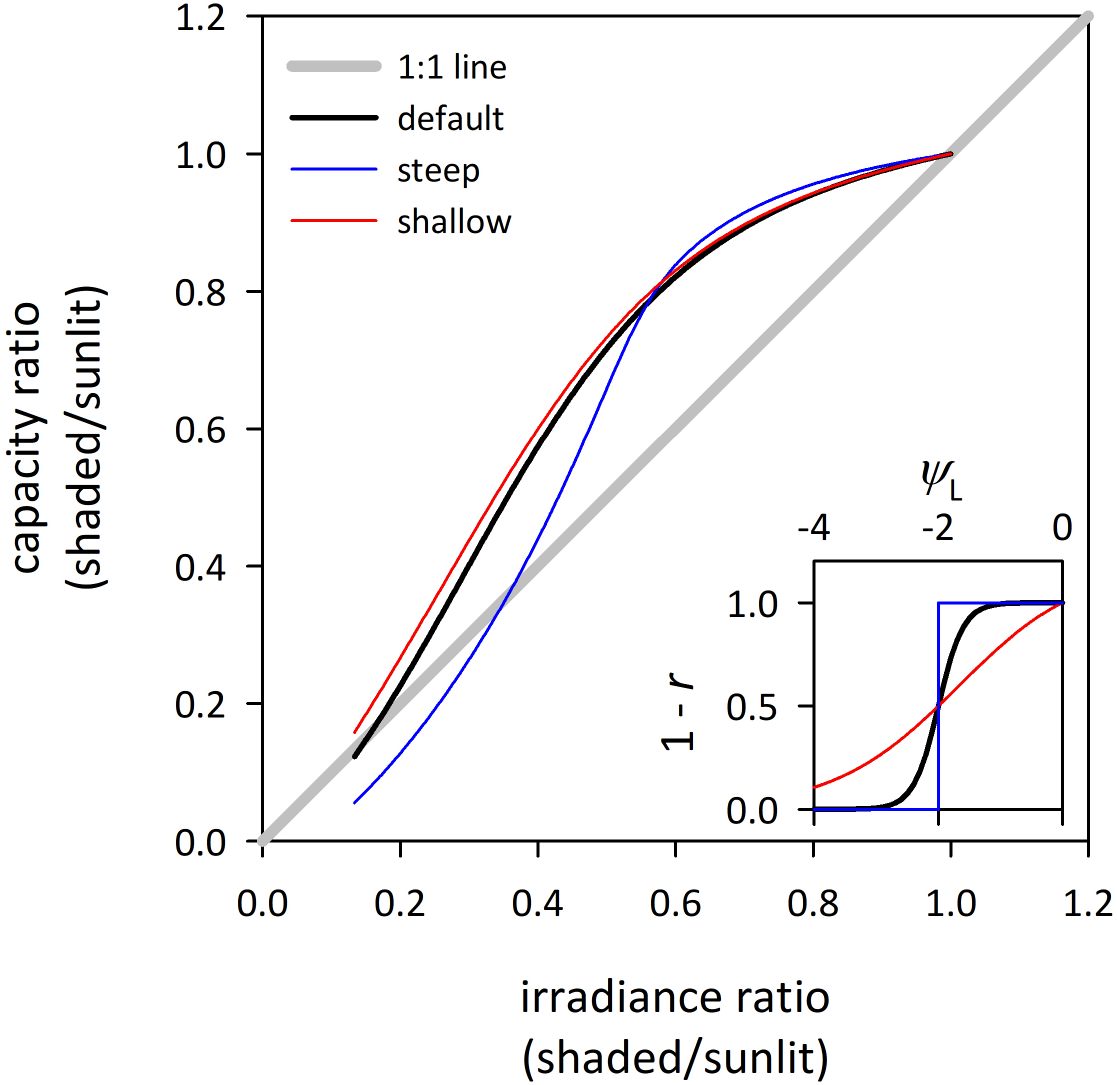
The divergence of the ratio of photosynthetic capacity between canopy modules from the irradiance ratio was greater if the risk function (*r*) used to penalize low water potentials (Eqn 4, shown inset as 1 – *r* vs *ψ*_L_) was less steep (red lines; *ξ* = 1 MPa^−1^), and conversely, the divergence was smaller if the risk function was infinitely steep (blue lines; *ξ* → ∞), relative to the default simulation (black lines;*ξ* = 5 MPa^−1^).

The patterns of photosynthetic capacity vs irradiance described above also gave rise to systematic differences in other parameters between the two modules. For example, intercellular CO_2_ concentration (*c*_i_) was greater in the shaded module than in the sunlit module, and the ratio of *c*_i_ between the two modules increased as the ratio of irradiance decreased (Fig 5). Similarly, the predicted optimal values of the marginal carbon product of water (∂*A*/∂*E*) and hydraulic conductance (∂*A*/∂*K* = [∂*A*/∂*E*]·[∂*E*/∂*K*] = [∂*A*/∂*E*]·[*ψ*_soil_ – *ψ*_L_]) were smaller in the shaded module than in the sunlit module; ∂*A*/∂*K* decreased more steeply in the canopy than ∂*A*/∂*E* because *ψ*_L_ was less negative in shaded modules than in sunlit modules (Fig 5). The value of the marginal carbon product of nitrogen (∂*A*/∂*N*) differed slightly between the two modules, but the marginal effect of N on carbon profit (∂*P*/∂*N*) was always equal between the two modules (Fig 5).

**Figure 5.**
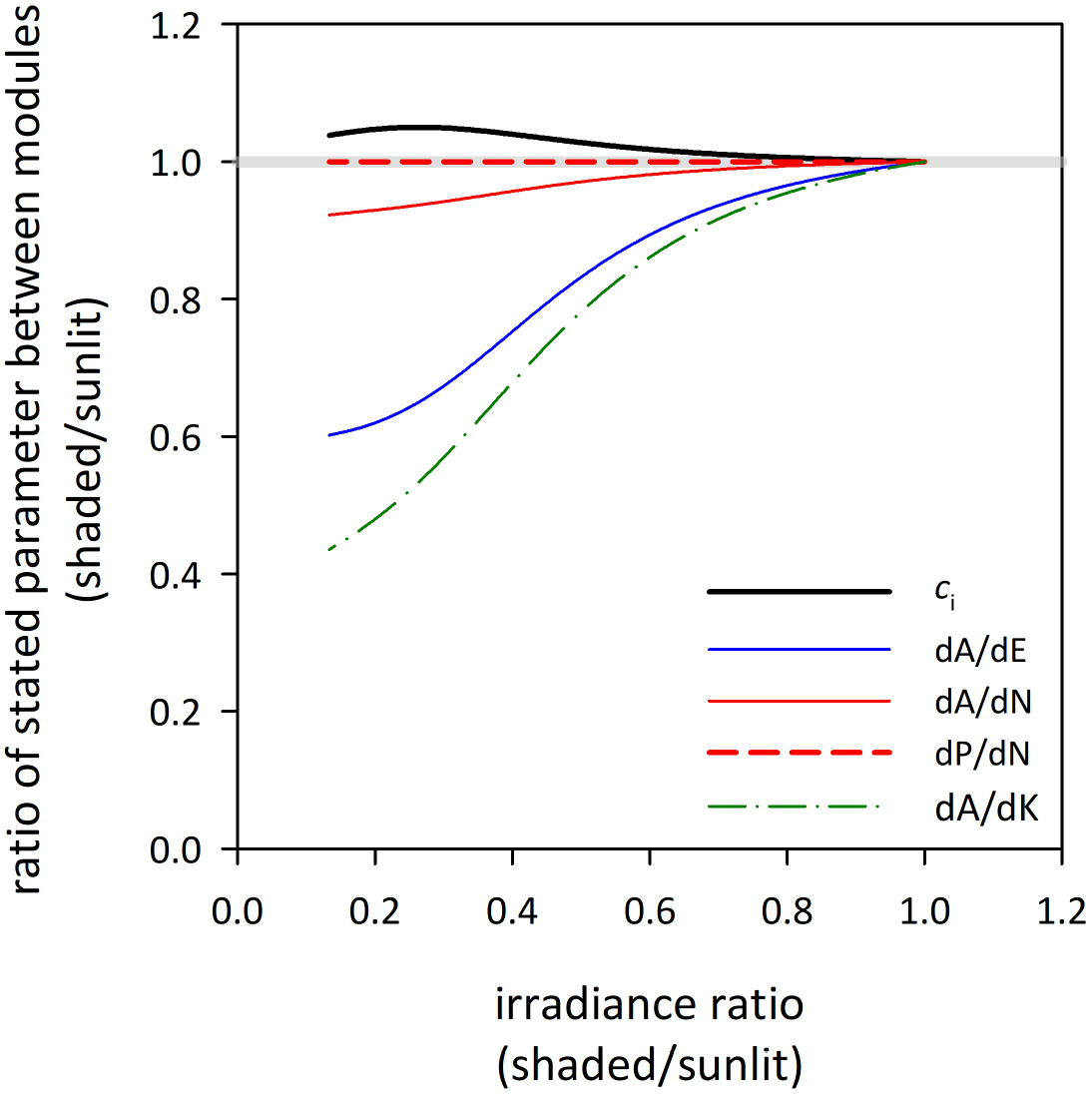
Other parameters of gas exchange differed systematically between canopy modules when carbon partitioning was adjusted among roots and stem C pools in both modules so as to maximize whole-plant carbon profit. Intercellular CO_2_ concentration (*c*_i_, black line) was greater in the shaded module; the marginal carbon revenues of water (∂*A*/∂*E*, blue line) and hydraulic conductance (∂*A*/∂*K*, green dash-dot line) were both smaller in the shaded module, and the marginal carbon revenue of nitrogen (∂*A*/∂*N*, solid red line) was slightly smaller in the shaded module. However, the marginal carbon profit of nitrogen (∂*P*/∂*N*, dashed red line), which accounts for the carbon cost of nocturnal leaf respiration, was invariant through the canopy.

All results described above used whole plant carbon profit as the goal function for optimization of C and N partitioning and adjustment of leaf water potential. However, very similar patterns were predicted (with slightly reduced divergence of the capacity and irradiance ratios) if the goal function was instead taken as total canopy photosynthesis (the sum of contributions from the two modules) instead of carbon profit (Fig S1). In those simulations, it also emerged that ∂*A*/∂*N* was invariant among modules in the optimum.

Predictions from the simulations described above were broadly consistent with published experimental observations of the relationship between capacity and irradiance ratios within canopies, except for irradiance ratios below around 0.4 (Fig 6).

**Figure 6.**
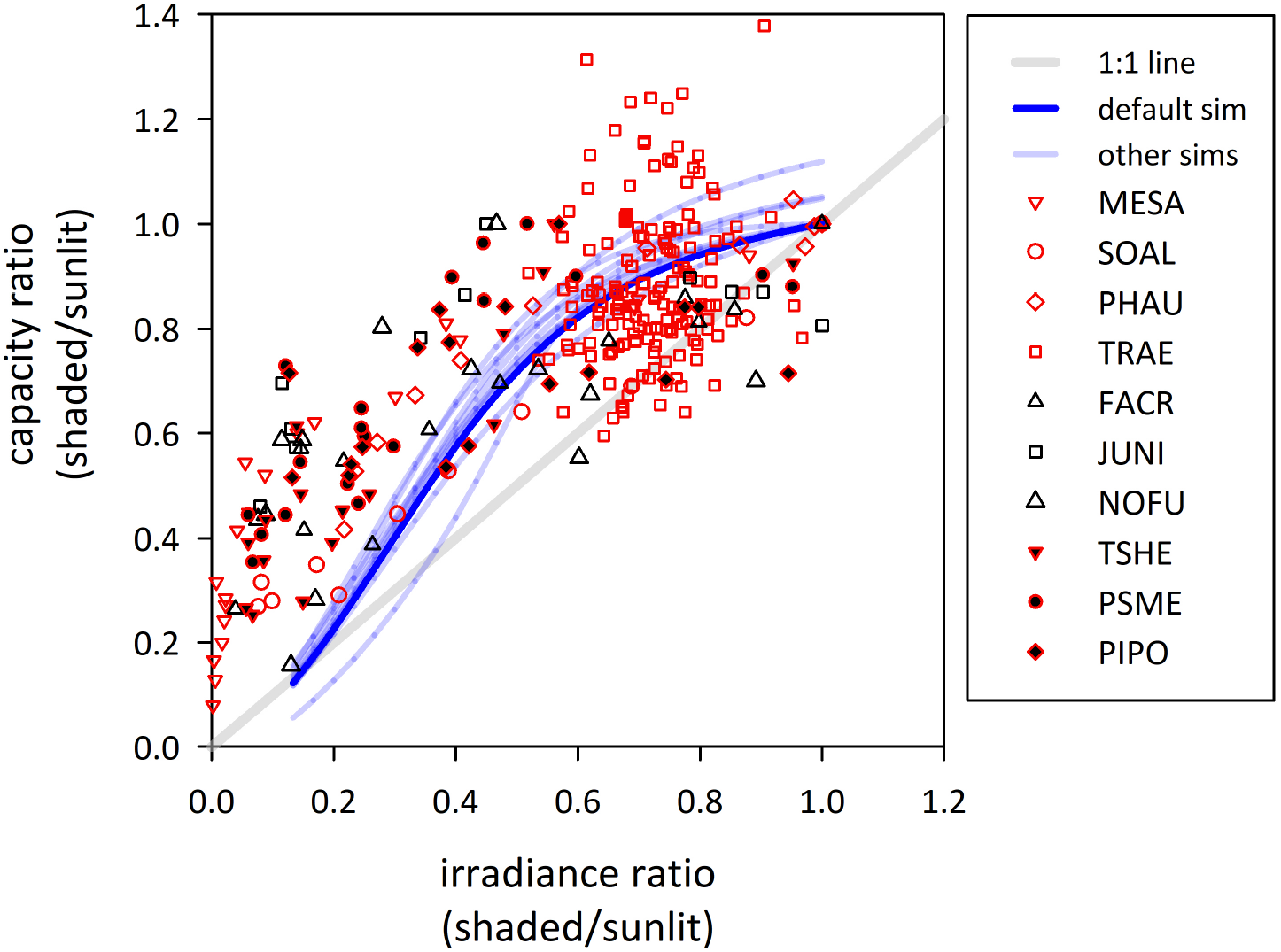
For large and intermediate ratios of incident irradiance between shaded and sunlit canopy modules, the simulations presented in this study (blue lines) are broadly consistent with experimental measurements from a range of species and environments (symbols). The solid blue line shows the default simulation from the present study; the dashed blue lines are the other simulations, reprinted from Figs 1, 2b, 3 and 4. Symbol coloring indicates species type (open red: herbaceous angiosperm; open black: woody angiosperm; closed red/black: woody gymnosperm). Species codes in the legend are as follows: MESA (*Medicago sativa*) (Louarn *et al.*, 2015), SOAL (*Solidago altissima*) (Hirose & Werger, 1987; Hirose *et al.*, 1988), PHAU (*Phragmites australia*) (Hirose & Werger, 1994, 1995), TRAE (*Triticum aestivum*) (Salter *et al.*, 2020), FACR (*Fagus crenata*) (Iio *et al.*, 2005), JUNI (*Juglans nigra x regia*) (Frak *et al.*, 2002), NOFU (*Nothofagus fuscata*) (Hollinger, 1996), TSHE (*Tsuga heterophylla*) (Bond *et al.*, 1999), PSMA (*Pseudotsuga menziesii*) (Bond *et al.*, 1999), PIPO (*Pinus ponderosa*) (Bond *et al.*, 1999). For MESA, capacity = electron transport capacity; for TRAE, capacity = light and CO_2_-saturated assimilation rate; for all other species, capacity = light-saturated assimilation rate at ambient CO_2_. Data for FACR, SOAL and PHAU were reproduced from Niinimets et al (2015).

## Discussion

Canopy profiles of photosynthetic capacity predicted by optimization theory have long been thought to contradict observations: leaves in shaded locations are thought to have too much photosynthetic capacity relative to their light availability, and conversely, sunlit leaves have too little capacity (recently reviewed by Niinemets *et al.*, 2015; Hikosaka *et al.*, 2016). I extended the theory to encompass not only optimal N partitioning among and within leaves, but also optimal carbon partitioning among functional C pools in the plant, while accounting for constraints on leaf water potential imposed by the risk of catastrophic desiccation. The patterns of within-canopy gas exchange that emerged from the resulting optimization are broadly similar to observations (Fig 6). This result suggests that optimization theory does not in fact contradict observations regarding canopy profiles of gas exchange parameters.

### Why does optimal carbon partitioning lead to variation in ∂*A*/∂*E* within the canopy?

I found that it is not optimal for ∂*A*/∂*E* to be invariant within a canopy, contrary to my own previous assertions (Buckley *et al.*, 2002, 2014; Farquhar *et al.*, 2002). To understand why, it helps to examine the Lagrange multiplier approach to optimization, which gave rise to my earlier assertions. In that approach, the optimum is found by setting the derivative of total photosynthesis with respect to stomatal conductance, *g*_sw_, equal to zero. The total supply of water – the total transpiration rate – is considered constant. Because the supply is constant, its derivative with respect to *g*_sw_ is zero, so it can be multiplied by an arbitrary constant (the Lagrange multiplier) and subtracted from photosynthesis without affecting the location of the optimum.

An often-tacit assumption of this approach is that the resource can be redistributed arbitrarily. That is, any leaf can have any transpiration rate, as long as the total for all leaves adds up to some imposed constant. However, that assumption is not valid for canopy transpiration, because water use by canopy elements is constrained by factors that cannot themselves vary arbitrarily. These include leaf water potential (*ψ*_L_), which is limited by fundamental biophysical constraints, and by root, stem and leaf hydraulic conductances, which depend on carbon investments. The transpiration rate *E* of a canopy module is approximately

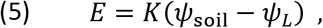

where *ψ*_soil_ is soil water potential and *K* is the total hydraulic conductance between the soil and the canopy module, including contributions from the module’s own stem tissues. Neither *K* nor *ψ*_L_ can vary arbitrarily as needed to allow transpiration rate to take on any possible value in any given leaf. Leaf water potential is not free to vary arbitrarily for two reasons. Firstly, it is constrained to remain above a critical threshold, *ψ*_c_, below which runaway loss of hydraulic conductivity becomes certain (Tyree & Sperry, 1989). Any strategy that leads to certain death is obviously inconsistent with the premise of optimization theory, so we must adopt the constraint that *ψ*_L_ ≥ *ψ*_c_. Secondly, economics also constrains how water potential can vary above that threshold. Imagine a canopy module in which *ψ*_L_ were always well above *ψ*_c_ (even allowing for a “safety margin”, discussed later). If water potential were then allowed to decline somewhat while remaining safely above *ψ*_c_, the rate of water transport to that module – and hence the transpiration rate, stomatal conductance and assimilation rate – would increase, without any cost to the plant. Therefore, having leaf water potential remain permanently above a safe lower limit is inherently suboptimal with respect to carbon gain.

Two counterarguments arise from game-theoretical considerations, but neither contradicts the arguments above. Firstly, it could be beneficial for a plant to keep *ψ*_L_ high to conserve water under some circumstances (for example, “banking” soil water stores for later in the season, or to mitigate the risk of fatal desiccation in droughts of uncertain duration) (Cowan, 1982; Mäkelä *et al.*, 1996; Lu *et al.*, 2016). Yet, for such a plant to make full use of its carbon investments in water uptake and transport, it must nevertheless allow *ψ*_L_ to approach a safe lower limit at some time, and throughout the canopy – at which point biophysics and economics would constrain *ψ*_L_ from below and above, respectively, as described earlier. Secondly, real leaves need to maintain a “safety margin” between *ψ*_L_ and *ψ*_c_, to prevent rapid excursions in evaporative demand or soil water potential from causing fatal excursions of *ψ*_L_ below *ψ*_c_ (Sperry, 2000; Delzon & Cochard, 2014). But that need also reflects a biophysical constraint: if stomata could respond instantly to the environment, transient excursions would be impossible and the safety margin would reduce to an infinitesimal sliver. Finite stomatal response rates are biophysical constraints like *ψ*_c_ itself, so the size of the safety margin is dictated by biophysical constraints (on stomatal kinetics) and environmental variables (the likelihood of dangerously rapid excursions in evaporative demand or soil water potential) (Meinzer *et al.*, 2017), and therefore does not represent freedom for leaf water potential to vary arbitrarily as needed to satisfy Eqn 5.

If *ψ*_L_ is not a truly free parameter, the only remaining parameter in Eqn 5 that the plant can control is *K*, which the plant can adjust via carbon partitioning. It follows that the optimal spatial distribution of water use (and by extension the total water use across the canopy) is defined by the optimal pattern of carbon partitioning. That is, canopy transpiration rate is an outcome of optimization, so it cannot be treated as a prior constraint in the optimization problem. The solution that arises from treating canopy transpiration as an imposed constant – that ∂*A*/∂*E* should be spatially invariant – is therefore invalid.

### Why is it optimal for ∂*A*/∂*E* and ∂*A*/∂*K* to be smaller, and photosynthetic capacity per unit irradiance greater, in shaded leaves?

The argument presented earlier explains why the Lagrange multiplier approach is inappropriate for identifying the optimal spatial distribution of water loss in the canopy, but it does not explain why ∂*A*/∂*E* should vary in the specific manner predicted. A simple thought experiment can explain this result. First, it stands to reason that hydraulic conductance should be smaller in shaded modules than in sunlit modules, because the rates of photosynthesis and transpiration are smaller in shaded modules. For a given module leaf area, the only way to achieve smaller module *K* is by reducing carbon investment in the module’s stem component, which determines stem hydraulic conductance (*K*_S_). However, because increasing investment in stem carbon earns diminishing returns in terms of module water use (that is, ∂^2^*E*/∂*C*_S_^2^ < 0; Eqn S7 in Supporting Information Notes S2), reducing *K*_S_ leads to an increase in the marginal return on stem carbon (∂*E*/∂*C*_S_). To reconcile that with optimal carbon partitioning – which requires the marginal sensitivity of carbon gain to stem carbon (∂*A*/∂*C*_S_ = [∂*A*/∂*E*]·[∂*E*/∂*C*_S_]) to be invariant (Buckley & Roberts, 2006a) – it follows that ∂*A*/∂*E* must be smaller in shaded modules than in sunlit modules.

This reasoning also resolves the apparent contradiction between my results and those of Peltoniemi et al. (2012), who concluded that photosynthetic capacity and irradiance should remain proportional between shaded and sunlit leaves if both hydraulic conductance and nitrogen are distributed optimally. Peltoniemi et al. (2012) assumed that optimal distribution of hydraulic conductance is equivalent to invariance in the sensitivity of assimilation rate to *K* (∂*A*/∂*K* ≈ [∂*A*/∂*E*]·[∂*E*/∂*K*] ≈ [∂*A*/∂*E*]·(*ψ*_soil_ – *ψ*_L_)), but as discussed earlier and illustrated in Fig 4, optimal carbon partitioning actually requires ∂*A*/∂*K* to vary between canopy modules. My simulations suggest that, in practice, differences in both ∂*A*/∂*E* and *ψ*_L_ contribute to satisfying this requirement. If *ψ*_L_ has less room to vary (due to a very steep risk curve), then ∂*A*/∂*E* must differ more between modules, and vice versa (Fig 4).

Other factors that commonly differ between shaded and sunlit modules, such as evaporative demand (Δ*w*) and hydraulic pathlength (*l*), can magnify differences in ∂*K*/∂*C*_S_ and/or ∂*A*/∂*K*, and thus also in photosynthetic capacity per unit irradiance. For example, if water must travel farther to reach sunlit leaves, then ∂*K*/∂*C*_S_ will be smaller for sunlit modules for a given KS (Eqn S5 in Supporting Information Notes S2); invariance of ∂*A*/∂*C*_S_ then requires either even-larger ∂*A*/∂*E* or even-lower *ψ*_L_ in sunlit leaves. My results suggest the relative importance of Δ*w* and pathlength in driving variation in capacity per unit irradiance may vary widely with conditions (Figs 2&3), consistent with a recent study (Bachofen *et al.*, 2020). Importantly, however, differences in evaporative demand and pathlength are not required to explain the general observation that shaded leaves have more N relative to their light environments: that pattern emerges as optimal even if Δ*w* and pathlength are identical between modules (Figs 2&3).

### Why does optimal carbon partitioning <u>not</u> lead to variation in ∂*A*/∂*N* within the canopy?

I found, as previously suggested (e.g., Field, 1983; Eqn 1), that it is optimal for the marginal carbon product of nitrogen, ∂*A*/∂*N*, to be invariant in a canopy (if the goal function is carbon profit [P] rather than canopy photosynthesis, then it is ∂*P*/∂*N* that must be invariant; Fig 4). Why does the argument developed earlier in relation to ∂*A*/∂*E* not apply to nitrogen? The reason is that nitrogen can, in principle, be partitioned arbitrarily between canopy modules. Although any given N partitioning may not be economically sensible – for example, building a high-N leaf in the shade would be uneconomical – there is no obvious biophysical constraint coupling C and N partitioning among modules, as there is for water. The total supply of photosynthetic N for the canopy can therefore be treated as a prior constraint for identifying optimal N distributions, so Eqn 1 remains valid.

### Implications for predicting and interpreting canopy profiles of gas exchange parameters

I found that it is generally optimal for photosynthetic capacity per unit irradiance to be greater in shaded leaves than in sunlit leaves. This has two significant implications. Firstly, it suggests that optimal N distribution does not generally make leaf-scale models of photosynthesis scale-invariant (Farquhar, 1989). The principle that optimization implies scale-invariance is the basis for “big-leaf” scaling of photosynthesis from leaves to canopies. However, efficient and accurate scaling procedures exist that can accommodate empirical profiles of N (e.g., de Pury & Farquhar, 1997), and at any rate, it has long been understood that actual canopy profiles diverge from those assumed in big-leaf models, so the failure of scale-invariance has few if any practical implications for modeling. Secondly, observed canopy profiles of photosynthetic capacity are not necessarily suboptimal, as has long been suspected. Recent work has suggested that genetic variation in these profiles could be used to improve canopy carbon gain in crops (Townsend *et al.*, 2018; Yin *et al.*, 2019; Salter *et al.*, 2020). My results do not necessarily contradict that idea, but they do suggest that “optimal” profiles are not necessarily those in which the capacity and irradiance ratios closely track one another through the canopy. More generally, by reconciling optimization theory with one class of observations that have long been thought to contradict the theory, my results support the use of optimization to predict plant form and function.

### Conclusions

Optimization of carbon partitioning in a whole plant model predicts that it is optimal for photosynthetic capacity per unit irradiance to be greater in more shaded modules than in more sunlit modules, thus helping to reconcile optimization theory with observations. This result holds in the absence of differences in evaporative demand or hydraulic pathlength between modules. Spatial invariance of the marginal carbon revenue of nitrogen, ∂*A*/∂*N*, is optimal as previously noted, but the marginal carbon revenues of both water (∂*A*/∂*E*) and hydraulic conductance (∂*A*/∂*K*) should vary through the canopy in the optimum.

## Acknowledgements

I thank the National Science Foundation (Awards #1557906 and #1951244) and the USDA National Institute of Food and Agriculture (Hatch project 1016439 and Award #2020-67013-30913) for support.

## Appendix

## Photosynthesis and carbon profit

The net photosynthesis rate for a given module (*A*_(m)_) is calculated from the FvCB model (Farquhar *et al.*, 1980), assuming that photosynthesis can be limited either by RuBP carboxylation (*A*_(m)_ = *A*_v(m)_) or by RuBP regeneration (*A*_(m)_ = *A*_J(m)_):

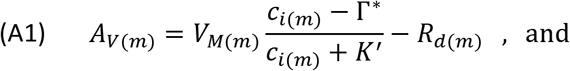

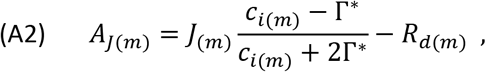

where *V*_M(m)_ is carboxylation capacity, *J*_(m)_ is potential electron transport rate, *c*_i(m)_ is intercellular CO_2_ concentration, Γ∗ is photorespiratory CO_2_ compensation point, *K*’ is the effective Michaelis constant for carboxylation, and *R*_d(m)_ is the rate of non-photorespiratory CO_2_ release in the light. *J* is computed as the hyperbolic minimum of the maximum potential electron transport rate (*J*_M(m)_) and the product of effective quantum yield of electrons (*ϕ*) and incident PPFD (*i*_(m)_):

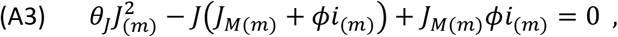

where *θ*_J_ is a dimensionless convexity parameter ≤ 1. *ϕ* is given by *ϕ* = 0.5·α·*ϕ*_PSIImax_, where *ϕ*_PSIImax_ is the maximum quantum yield of photosystem II, and *α* is leaf absorptance to photosynthetically active radiation, which depends on chlorophyll content (Chl, mmol m^−2^) (Evans, 1996):

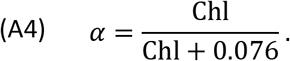

Chl depends on N invested in light capture (*N*_C(m)_) and electron transport (*N*_J(m)_) as (Buckley *et al.*, 2013)

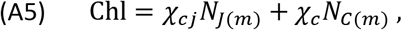

*V*_M(m)_ and *J*_M(m)_ depend on module-wise N pools for Rubisco (*N*_v(m)_) and electron transport:

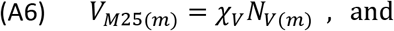

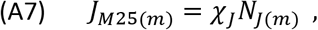

where χ_v_ and χ_J_ are fixed parameters and the subscripts “25” indicate values at 25°C. The parameters *V*_M_, *J*_M_, Γ∗, K’, *ϕ*_PSIImax_, *θ*_J_ and *R*_d_ all depend directly on temperature (see Supporting Information Notes S3). I assumed that *R*_d(m)_ is proportional to *V*_M(m)_ at 25°C, such that *R*_d25(m)_ = 0.0089·*V*_M25(m)_ (de Pury & Farquhar, 1997)‥

Intercellular CO_2_ concentration is determined by the balance between CO_2_ demand by the mesophyll (Eqns A1 and A2) and diffusional supply through the stomata:

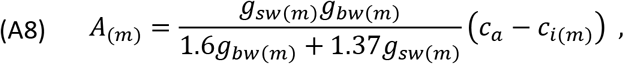

where *c*_a_ is ambient CO_2_ concentration, and *g*_sw_ and *g*_bw_ are stomatal and boundary layer conductances to water vapor, respectively (Eqn A8 ignores mesophyll conductance). Combining Eqn A8 with either Eqn A1 or A2 produces a quadratic expression for *c*_i(m)_, whose solution for *c*_i_ can be applied to Eqn A1 or A2 to determine *A*_v(m)_ or *A*_J(m)_, respectively. Assimilation rate is usually calculated as the simple minimum of *A*_v(m)_ and *A*_J(m)_; because that produces discontinuities in the sensitivity of A to *N*_v(m)_, *N*_J(m)_, and *g*_sw(m)_, which can preclude unambiguous identification of optima, I “smoothed” the transition between *A*_v(m)_ and *A*_J(m)_ by computing *A*_(m)_ as the hyperbolic minimum of *A*_v(m)_ and *A*_J(m)_:

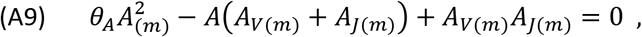

and set the dimensionless parameter *θ*_A_ to 0.999. Canopy net carbon gain is (*A*_c_) the sum of the products of leaf area (*L*_(m)_) and photosynthesis per unit leaf area (*A*_(m)_) for each canopy module *m* (*m* = 1 or 2 in this study):

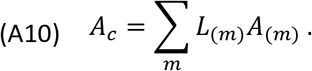

I modeled carbon profit (*P*) as

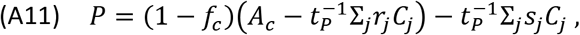

where *f*_c_ is the fraction of net allocatable carbon (i.e., net photosynthesis minus whole-plant maintenance respiration) lost to construction respiration when constructing the carbon pools, *t*_P_ is the effective number of seconds per year of active photosynthesis (which scales instantaneous photosynthesis rates to annual values), and *r*_j_ and *s*_j_ are the maintenance respiration rate and senescence rate per unit carbon for carbon pool *j*, respectively. I calculated *s*_j_ as the inverse of tissue lifespan, *τ*_j_. Note that *r*_j_ for leaf carbon pools represents nocturnal maintenance respiration (the photosynthesis model described above includes daytime maintenance respiration as *R*_d_); thus *r*_L(m)_·*C*_L(m)_/*t*_p_ = *L*_(m)_·*R*_n(m)_, where *R*_n(m)_ is the nocturnal leaf respiration rate (estimated as *R*_d(m)_ divided by the proportion by which the instantaneous value of leaf respiration is inhibited in the light below the value in the dark).

Canopy photosynthesis is largely determined by parameters that depend on three resources, or *inputs*: light (*i*), water (which limits *g*_sw_), and nitrogen (which limits *V*_M_, *J*_M_ and *α*). The amount of each resource available to a given module depends on the availability of those resources in the environment, but also on the sizes of functional carbon pools (roots, stems and leaves) that acquire and transport those resources. The next section describes models for those dependencies.

## Supply of photosynthetic nitrogen to the canopy

I modeled the total supply of photosynthetic N available for partitioning between canopy modules (*N*_L_) by assuming a steady-state between N uptake by roots (*U*_N_) and losses due to tissue senescence. I modeled *U*_N_ following Buckley and Roberts (Buckley & Roberts, 2006b), as

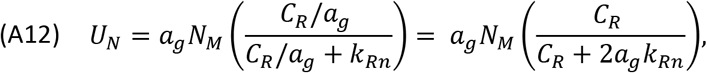

where *a*_g_ is ground area, *N*_M_ is the rate of N inputs into the soil per unit ground area (mmol N m^−2^_ground_ s^−1^), *C*_R_ is root carbon, *k*_Rn_ is the value of root C (per unit ground area) at which the rate of N uptake is half of *N*_M_. I set *a*_g_ to twice the projected ground area of each canopy module, because the root system as modeled here supplies both modules. I modeled the N loss rate to leaf senescence as

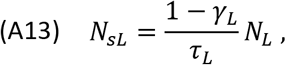

where *τ*_L_is leaf lifespan (s) and *γ*_L_ is the fraction of N withdrawn from leaves prior to senescence. The N loss rates due to senescence of N-containing non-photosynthetic tissues (roots and stems) are

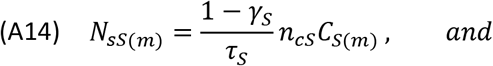

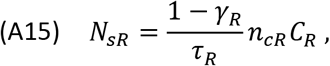

for stems and roots, respectively, where *γ*_j_, *τ*_j_ and *N*_cj_ are the fractions of N withdrawn from pool j before senescence, the lifespan of the pool, and the N:C ratio of the pool, respectively. Setting *U*_N_ – *N*_sL_ – *N*_sS_ – *N*_sR_ equal to zero and solving for *N*_L_ gives

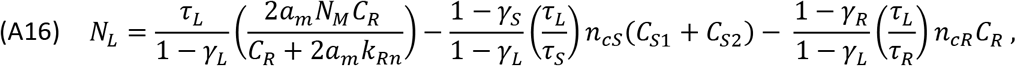

where *C*_S1_ and *C*_S2_ are stem carbon in canopy modules #1 and #2, respectively.

## Water uptake and transport, and transpiration and stomatal conductance

At steady-state and on average, mass conservation requires that the total transpiration rate of a module (*E*_t(m)_) equals the rate of water transport from the soil to the module. The latter rate is determined by hydraulic conductances and soil and leaf water potentials. Because both modules share the root component of whole-plant hydraulic conductance (*K*_R_), *E*_t_ in each module depends in part on the leaf water potential and total hydraulic conductance of the other module. The resulting expressions for *E*_t1_ and *E*_t2_, derived in Supporting Information Notes S4, are

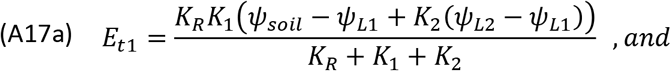

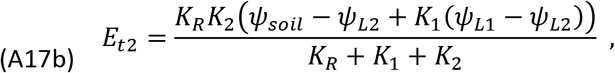

in which *K*_1_ = (*K*_S1_^−1^ + *K*_L1_^−1^)^−1^ and *K*_2_ = (*K*_S2_^−1^ + *K*_L2_^−1^)^−1^, where *K*_S(m)_ and *K*_L(m)_ are the stem and leaf hydraulic conductances of module m, respectively. The hydraulic conductances depend on carbon in each pool:

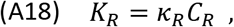

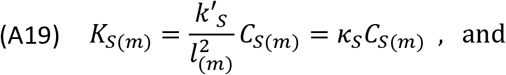

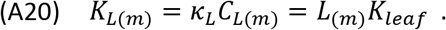

Equations A18–A20 are derived in Supporting Information Notes S5; *κ*_R_, k’_S_ and *κ*_L_ are parameters that I treated as constants in this study, and *l*(_m_) is the hydraulic pathlength of canopy stem module *m*. The module transpiration rate must also equal the product of module leaf area (*L*(_m)_), total conductance to water vapor, and leaf to air water vapor mole fraction difference (Δ*w*(_m)_):

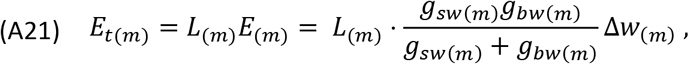

where *g*_sw_ and *g*_bw_ are stomatal and boundary layer conductances to H_2_O, respectively (note *g*_sc_ = *g*_sw_/1.6 and *g*_bc_ = *g*_bw_/1.37). Δ*w* depends on leaf temperature, *T*_L_:

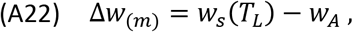

where *w*_A_ is the ambient water vapor mole fraction and *w*_s_(*T*) is the saturated water vapor mole fraction at a temperature *T* (*w*_s_(*T*) = 611.2·exp(17.62·*T*/(243.12 + *T*))/*P*_A_, where *P*_A_ is atmospheric pressure in Pa and *w*_s_ is in mol mol^−1^ (World Meteorological Organization, 2008)). *w*_A_ = RH·*w*_s_(*T*_A_), where RH is relative humidity as a fraction. I estimated *T*_L_ based on energy balance (Eqn A23, derived in Supporting Information Notes S6):

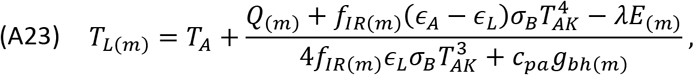

where *T*_A_ is air temperature in deg C (*T*_AK_ in kelvins), *Q* is absorbed shortwave radiation, *E* is leaf transpiration rate (calculated from Eqn A17 and module leaf area), *g*_bh_ is boundary layer conductance to heat (= *g*_bw_/1.08), *ε*_A_ is the atmospheric emissivity (*ε*_A_ = 0.642·(*P*_A_*w*_A_/*T*_AK_)^1/7^ (Leuning *et al.*, 1995)), *ϕ*_L_ is leaf emissivity (0.97), *λ* is the latent heat of vaporization (4.4·10^4^ J mol^−1^), σ_B_ is the Stefan-Boltzmann constant (5.67·10^−8^ J m^−2^ s^−1^ K^−4^) and *c*_pa_ is the heat capacity of air (29.2 J mol^−1^). *f*_IR_ is the leaf-atmosphere IR exchange as a fraction of the value at the top of the canopy. I estimated *f*_IR_ by assuming total visible and IR radiation were attenuated through the canopy with extinction coefficients *k*_i_ and *k*_d_, respectively, giving *f*_IR_ = (*i/i*_top_)^kd/ki^, where *i*_top_ is the incident irradiance above the canopy (= *i*_sunlit_). Additional details regarding Eqn A23 are given in the SI.

The stomatal conductance in each module (*g*_sw(m)_) is found by inverting Eqn A21:

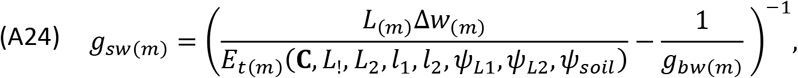

where *E*_t(m)_ is calculated from Eqns A17–A20 based on the vector of carbon pools (C), the axial stem lengths (*l*_1_ and *l*_2_) and water potentials (*ψ*_L1_, *ψ*_L2_) of both modules, and *ψ*_soil_. Finally, *g*_sw(m)_ is appliedto Eqn A8 to calculate CO_2_ assimilation rate.

## Degrees of freedom

The model for canopy photosynthesis and carbon profit has nine biological degrees of freedom (dfs), excluding parameters treated as constants. There are 15 variables: *N*_v(m)_, *N*_J(m)_, *N*_C(m)_, *i*_(m)_, *g*_sw(m)_, *ψ*_L(m)_, and *C*_S(m)_, each for two modules, plus *C*_R_. Two dfs are removed by specifying incident irradiance for each module; one df is removed by constraining the sum of the remaining C pools to a constant (*C*_S1_ + *C*_S2_ + *C*_R_ = *C*_T_); Eqn A24 removes two dfs by defining *g*_sw(m)_ as a function of other variables among the 17 listed above; and Eqn A16 removes one df by constraining the sum of photosynthetic N pools (*L*_1_(*N*_V1_ + *N*_J1_ + *N*_C1_) +*L*_2_(*N*_V2_ + *N*_J2_ + *N*_C2_) = *N*_L_). Of the nine remaining dfs, seven can be expressed as partitioning fractions (two for C and five for N), and two represent the leaf water potential in each module.

