## Supporting Information File S2 for "Optimal carbon partitioning reconciles the apparent divergence between optimal and observed canopy profiles of photosynthetic capacity"

Contents:


- ***Notes S1. Parameter estimation***
- ***Figure S1. Simulations with carbon gain rather than carbon profit as the goal.***
- ***Notes S2. Response of water use to stem carbon investment, and comparison between modules***
- ***Notes S3. Temperature dependencies of photosynthetic parameters***
- ***Notes S4. Derivation of expressions for water use in each module***
- ***Notes S5. Derivation of expressions for hydraulic conductances***
- ***Notes S6. Derivation of expression for leaf temperature***


*(Note: Supporting Information File S1 contains the Excel file used to perform the simulations described in the paper.)*

---

**Notes S1. Parameter estimation**

I estimated the values of several parameters in the model from the literature, as described below.

a. $\boldsymbol{f_c}$ (the fraction of net carbon gain used in construction respiration). I estimated fc from data given by Poorter and de Jong (1999), who reported median construction costs of 1.47 g glucose $g^{-1}$ dry mass and tissue carbon content of 2.19 g dry mass $g^{-1}$ carbon. Given 180 g glucose $mol^{-1}$ glucose, 6 mol C $mol^{-1}$ glucose and 12 g C $mol^{-1}$ C, this gives construction costs of 1.286 g C $g^{-1}$ C. Thus, $f_c = 0.286$.

b. $\boldsymbol{\gamma_L}$**,** $\boldsymbol{\gamma_S}$ (fraction of N withdrawn from leaf and stem tissues, respectively, prior to senescence). I used a value of 0.5 for both $\gamma_L$ and $\gamma_S$ (Murty *et al.*, 1996), the latter based on the ratio of N:C in senescing woody litter vs new wood.

c. $\boldsymbol{k_i}$ (effective extinction coefficient for visible light). I used $k_i = 0.56$, the mean across all biomes reported by Zhang et al (2014) in a global meta-analysis of $k_i$.

d. $\boldsymbol{K_{leaf}}$ (leaf hydraulic conductance). I used the all-species average of $K_{leaf} = 11.46$ mmol $m^{-2}$ $s^{-1}$ $MPa^{-1}$ (0.01146 mol $m^{-2}$ $s^{-1}$ $MPa^{-1}$) reported by Sack and Holbrook (2006).

e. $\boldsymbol{\kappa_R}$ (hydraulic conductance per unit fine root carbon). I used the estimate given by Buckley and Roberts (2006), $\kappa_R = 6.6 \cdot 10^{-4}$ mol $s^{-1}$ $MPa^{-1}$ $mol^{-1}$, which was based on data from a 40-year old lodgepole pine stand (Ryan & Waring, 1992), and the assumption that in those trees, half of whole-plant resistance was in the root system.

f. $\boldsymbol{k_{RN}}$ (root carbon per unit ground area at which N uptake is half its maximum). I used the estimate given by Buckley and Roberts (2006), $k_{RN} = 16$ mol $m^{-2}$, which was based on data from a 40-year old lodgepole pine stand (Ryan & Waring, 1992), and the assumption that N uptake and losses were in balance in those trees and that N uptake was half-saturated by root C.

g. $\boldsymbol{k'_S}$ (effective sapwood permeability). From Eqn S22, $k'_S$ is given by

(S1) $$k'_S = \frac{k_S}{\eta V_w \zeta_S \rho_{cS}} ,$$

where $k_S$ is sapwood permeability, $\eta$ is the dynamic viscosity of water ($8.9 \cdot 10^{-10}$ MPa s at 25°C), $V_w$ is the molar volume of water ($1.8 \cdot 10^{-5}$ $m^3$ $mol^{-1}$), $\rho_{cS}$ is the carbon density of stem tissue, and $\zeta_S$ is a term that accounts for stem taper. I estimated $k_S$ from data presented by Sperry et al (2006) for a range of woody species, for which median $k_S$ was $1.38 \cdot 10^{-12}$ $m^2$ ($2.23 \cdot 10^{-12}$ and $6.54 \cdot 10^{-13}$ $m^2$ for vessels and tracheids, respectively). I estimated $\rho_{cS}$ from data given by Hacke et al (2001), who reported median values of 21192 and 25046 mol $m^{-3}$ for conifers and angiosperms, respectively; I used the mean of these two values (23119 mol $m^{-3}$). I used the value of $\zeta_S = 0.38$ given by Buckley and Roberts (2006), which implies the total volume of the stem component of a module is 38% of the volume of a cylinder with the same length and basal conducting area; that number was estimated from allometric data for lodgepole pine given by Reid et al. (1974) and Litton (2002). Together, this gives $k'_S$ at 25°C = 0.00979 mol $m^2$ $s^{-1}$ $MPa^{-1}$ $mol^{-1}$. Finally, to account for effects of temperature on viscosity (which decreases as the 7th power of absolute temperature), I multiplied this value by the ratio $(T_{AK}/298.15)^7$ in each simulation.

h. $\boldsymbol{n_{cR}}$**,** $\boldsymbol{n_{cS}}$ (N:C ratios of root and stem tissues, respectively). I used the value of $n_{cR}$ = 17 mmol N $mol^{-1}$ C given by Dewar and McMurtrie (1996) (this value happens to be equal to the global mean for roots < 1 mm in diameter given by Yuan et al. (2011)), and the value of $n_{cS}$ = 1.2 mmol $mol^{-1}$ given by Murty et al. (1996).

i. $\boldsymbol{r_R}$**,** $\boldsymbol{r_S}$ (maintenance respiration per unit C for roots and stems, respectively). I estimated $r_R$ from data given by George et al. (2003), who reported values of 0.68 and 0.76 g $g^{-1}$ $y^{-1}$ for loblolly pine and sweetgum trees, respectively; I used the mean of these two values (0.72). I estimated rS from data for sweetgum from Edwards et al. (2002), who reported annual stem maintenance respiration of 7.9 kg $CO_2$ $m^{-3}$ $y^{-1}$, which is equivalent to 0.0072 mol C $mol^{-1}$ C $y^{-1}$ using the stem carbon density value ($\rho_{cS}$ = 23119 mol $m^{-3}$) given above.

j. $\boldsymbol{\tau_S}$**,** $\boldsymbol{\tau_L}$**,** $\boldsymbol{\tau_R}$ (lifespans of stem, leaf and root tissues, respectively). I assumed leaf and fine root lifespans of 1 year. For the functional lifespan of conducting tissues in the stem, I took the average (9.27 y) of median values for six angiosperm species (9.43 y) and three gymnosperm species (9.1 y) collated from a range of studies (Bancalari *et al.*, 1987; Gebauer *et al.*, 2008; Wang *et al.*, 2010; van der Sande *et al.*, 2015).

k. $\boldsymbol{t_p}$ (time scaling factor that converts instantaneous mid-day rates calculated by the gas exchange model into annualized values). Several parameters in the model estimated from the literature were given in annualized units (e.g., mol C $y^{-1}$) that are not directly commensurable with the instantaneous units used in the photosynthesis model (e.g., $\mu$mol $s^{-1}$). Rigorous fusion of these two time scales would require explicit simulation of diurnal and seasonal timecourses, which would render the optimization procedure used in this study computationally infeasible. I therefore applied an approximate temporal scaling procedure, as follows. First, I defined the gas exchange rates (represented generically here as $Y$) simulated using the instantaneous leaf-level model described in the main text as mid-day maximum and seasonal peak values ($Y_{max,peak}$). Second, I assumed that daily maximum rates ($Y_{max}$) varied seasonally between zero and $Y_{max,peak}$, according to the function $Y_{max}(d) = Y_{max,peak} \cdot [1 - 0.5 \cdot \cos(2\pi \cdot d/365)]$, where $d$ is day of the year. The average of this function is $Y_{max,peak}/2$. Third, I assumed that diurnal rates varied sinusoidally between sunrise and sunset, peaking at $Y_{max}$, according to the function $Y(t) = Y_{max} \cdot \sin(\pi \cdot (t - t_{sunrise})/(t_{sunset} - t_{sunrise}))$ for $t$ (time) between sunrise ($t_{sunrise}$) and sunset ($t_{sunset}$), and $Y(t) = 0$ otherwise. The diel average of $Y(t)$ for a day with 12 hours of sunlight is then $Y_{max}/\pi$. Thus, the annual average instantaneous value of any given gas exchange rate, $\langle Y \rangle$, is approximately $Y_{max,peak}/2\pi$. Finally, converting from $\mu$mol $s^{-1}$ to mol $y^{-1}$ using $[3600\ s\ h^{-1}] \cdot [24\ h\ d^{-1}] \cdot [365\ d\ y^{-1}] \cdot [10^{-6}\ mol\ \mu mol^{-1}]$ gives $\langle Y \rangle/[mol\ y^{-1}] \approx 5 \cdot R_{max,peak}/[\mu mol\ s^{-1}]$ and $t_P = 5 \cdot 10^6\ s\ y^{-1}$.

---

**Figure S1**. Predicted optimal relationships between the ratios of photosynthetic capacity between shaded and sunlit modules (y-axis) and that of irradiance (x-axis), when the goal function for optimization was chosen as whole-plant carbon profit (red line) or canopy photosynthesis (blue line). Because the two results are very similar, the percent difference between them is plotted with a dash-dot green line (the horizontal shaded green line represents a % difference of zero.)

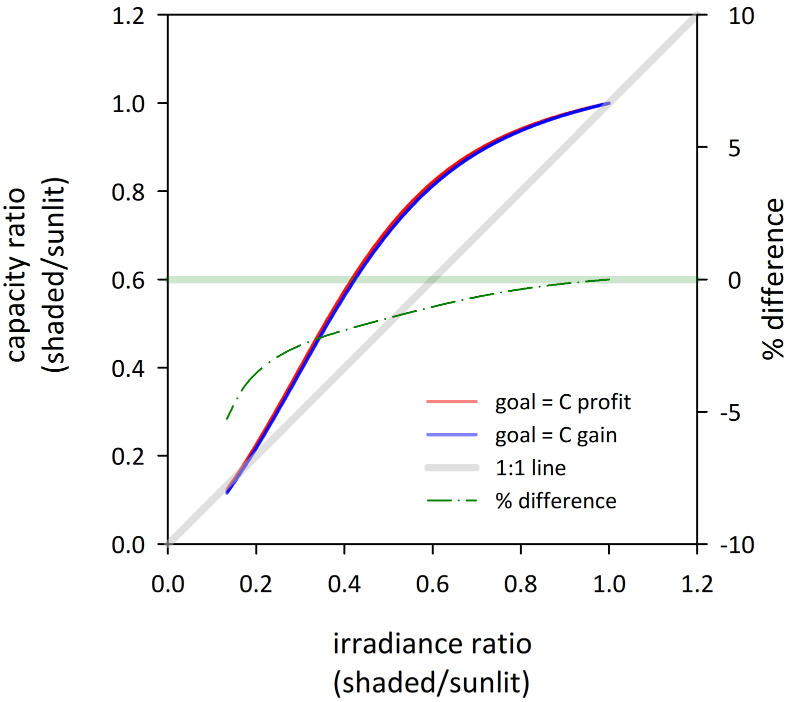

---

***Notes S2. Response of water use to stem carbon investment, and comparison between modules***

In the Discussion of the main text, I argued that the marginal return (in terms of water use) of stem carbon investment ($\partial E/\partial C_S$) will generally be larger in shaded canopy modules, with the result that the marginal carbon revenue of water ($\partial A/\partial E$) will be lower in shaded modules in order to satisfy the requirement that $\partial A/\partial C_S$ be identical between modules to optimize carbon partitioning. Here I demonstrate formally why this is so. The transpiration rate of a given module (#1) is

$$\text{(S2)} \qquad \frac{\partial E_{t1}}{\partial C_{S1}} = \frac{\partial E_{t1}}{\partial K_1}\frac{\partial K_1}{\partial K_{S1}}\frac{\partial K_{S1}}{\partial C_{S1}} \; .$$

From Eqn S29 in Notes S5 below, the first derivative on the right-hand side of is

$$\text{(S3)} \qquad \frac{\partial E_{t1}}{\partial K_1} = \frac{(K_R + K_2)\big(K_R(\psi_{soil} - \psi_{L1}) + K_2(\psi_{L2} - \psi_{L1})\big)}{(K_R + K_1 + K_2)^2} \; ;$$

from the definition of $K_1$ above Eqn S27, the second derivative in S2 is

$$\text{(S4)} \qquad \frac{\partial K_1}{\partial K_{S1}} = \left(\frac{K_{L1}}{K_{S1} + K_{L1}}\right)^2 \; ;$$

and from Eqn S22a, the third derivative in S2 is simply $\partial K_{S1}/\partial C_{S1} = k'_S/l_1^2$. Note that $\partial K_1/\partial C_{S1}$ is then

$$\text{(S5)} \qquad \frac{\partial K_1}{\partial C_{S1}} = \frac{k'_s}{l_1^2}\left(\frac{K_{L1}}{K_{S1} + K_{L1}}\right)^2 \; .$$

Combining these gives

$$\text{(S6)} \qquad \frac{\partial E_{t1}}{\partial C_{S1}} = \frac{k'_s}{l_1^2}\left(\frac{K_{L1}}{K_{S1} + K_{L1}}\right)^2 \frac{(K_R + K_2)\big(K_R(\psi_{soil} - \psi_{L1}) + K_2(\psi_{L2} - \psi_{L1})\big)}{(K_R + K_1 + K_2)^2} \; ,$$

This expression is a decreasing function of stem carbon for module #1 ($C_{S1}$ only affects terms in the denominator – $K_{S1}$ and $K_1$ – and both positively). Thus, it follows that investing stem carbon in a given module earns diminishing returns in terms of that module's transpiration rate; i.e.,

(S7) $$\frac{\partial^2 E_{t1}}{\partial C_{S1}^2} < 0 \, .$$

Maximizing canopy carbon gain requires that $\partial A_t/\partial C_{S1} = \partial A_t/\partial C_{S2}$, or

(S8) $$\frac{\partial A_{t1}}{\partial E_{t1}}\frac{\partial E_{t1}}{\partial C_{S1}} = \frac{\partial A_{t2}}{\partial E_{t2}}\frac{\partial E_{t2}}{\partial C_{S2}} \, ,$$

where $A_{t(m)}$ is the total assimilation rate of module *m* (the assimilation rate on a leaf area basis times the leaf area, which is equal by definition between the two modules in this study). Given that gas exchange is uniform within each module, $\partial A_t/\partial E_t = \partial A/\partial E$. Thus

(S9) $$\frac{\left(\frac{\partial A}{\partial E}\right)_2}{\left(\frac{\partial A}{\partial E}\right)_1} = \frac{\left(\frac{\partial E_{t1}}{\partial C_{S1}}\right)}{\left(\frac{\partial E_{t2}}{\partial C_{S2}}\right)} = \left(\frac{l_2}{l_1}\right)^2 \left(\frac{K_{S2}+K_L}{K_{S1}+K_L}\right)^2 \left(\frac{K_R+K_2}{K_R+K_1}\right)\left(\frac{\psi_{soil}-\psi_{L1}+\frac{K_2}{K_R}(\psi_{L2}-\psi_{L1})}{\psi_{soil}-\psi_{L2}+\frac{K_1}{K_R}(\psi_{L1}-\psi_{L2})}\right) ,$$

$K_{L1}$ and $K_{L2}$ have been replaced by a single value, $K_L$, as $K_{leaf}$ and module leaf area are assumed equal between modules in this study.

Now suppose module #2 is the "sunlit" module, and module #1 is the shaded module, so that in the optimum $K_{S2} > K_{S1}$ (and therefore $K_2 > K_1$) and $\psi_{L2} \leq \psi_{L1}$. Each of the ratios in parentheses on the right-hand side of Eqn S9 will then be greater than unity (unless the hydraulic pathlength for the shaded module is greater than that for the sunlit module – i.e., $l_1 > l_2$ – which is far less likely than the converse in real trees, as sunlit leaves tend to be located in more distal locations than shaded leaves, if anything). Therefore, the entire expression must be greater than unity, meaning that optimal carbon partitioning requires $\partial A/\partial E$ to be greater for module #2 (the sunlit module) than for module #1 (the shaded module).

---

***Notes S3. Responses of photosynthetic parameters to temperature***

Several parameters in the photosynthesis model presented in the main text depend on temperature. I used temperature responses given by Bernacchi et al (2001, 2003):

$$V_M = V_{M25} \cdot \exp\left(26.35 - \frac{65.33}{RT_{LK}}\right), \tag{S10}$$

$$J_M = J_{M25} \cdot \exp\left(17.7 - \frac{43.9}{RT_{LK}}\right), \tag{S11}$$

$$\Gamma_* = \exp\left(19.02 - \frac{37.83}{RT_{LK}}\right), \tag{S12}$$

$$K_C = \exp\left(38.05 - \frac{79.43}{RT_{LK}}\right), \tag{S13}$$

$$K_O = \exp\left(20.3 - \frac{36.38}{RT_{LK}}\right), \tag{S14}$$

$$\phi_{PSIImax} = 0.352 + 0.022T_L - 0.00034T_L^2, \tag{S15}$$

$$\theta_J = 0.76 + 0.018T_L - 0.00037T_L^2\ , \ \ and \tag{S16}$$

$$R_d = 0.0089V_{M25} \cdot \exp\left(18.72 - \frac{46.39}{RT_{LK}}\right), \tag{S17}$$

where $R$ = 0.00831446 and $V_{M25}$ and $J_{M25}$ are values at 25°C, calculated from nitrogen pools as described in the main text. The parameter $K'$ in the main text (the effective Michaelis constant for RuBP carboxylation) is equal to $K_C(1 + 210/K_O)$, which assumes oxygen is ~21% of the atmosphere by volume.

---

***Notes S4. Derivation of expressions for hydraulic conductances***

From Darcy's Law, the rate of water flow ($F_j$) through a tissue *j* with cross-sectional conducting area $S_j$ ($m^2$), permeability $k_j$ ($m^2$) and length $l_j$ (m) is

$$\text{(S18)} \quad F_j = \frac{k_j a_j}{l_j \eta V_w} \Delta P \ ,$$

where $\eta$ is dynamic viscosity of water ($1 \cdot 10^{-9}$ MPa s), $V_w$ is the molar volume of water ($1.8 \cdot 10^{-5}$ $m^3$ $mol^{-1}$) and $\Delta P$ is the axial pressure difference across the tissue. The hydraulic conductance, $K_j$ (flow per unit pressure or water potential gradient) is thus

$$\text{(S19)} \quad K_j = \frac{k_j S_j}{l_j \eta V_w} \ ,$$

To relate this to the carbon content of the tissue ($C_j$), note that the volume of conducting tissue ($V_j$) is

$$\text{(S20)} \quad V_j = \zeta_j S_j l_j \ ,$$

where $\zeta_j$<1 accounts for taper; thus $C_j$ is

$$\text{(S21)} \quad C_j = \rho_{cj} V_j = \rho_{cj} \zeta_j S_j l_j \ ,$$

where $\rho_{cj}$ is the carbon density of the tissue (mol $m^{-3}$). Thus, $S_j = C_j/(\rho_{cj} \cdot \zeta_j \cdot l_j)$. Applying this to Eqn S19 gives

$$\text{(S22a)} \quad K_j = \frac{k_j C_j}{l_j^2 \eta V_w \rho_{cj} \zeta_j} = \frac{k'_j}{l_j^2} C_j \ , \ \ or$$

$$\text{(S22b)} \quad K_j = \kappa_j C_j \ ,$$

where $k'_j = k_j/(\eta \cdot V_w \cdot \rho_{cj})$ and $k_j = k'_j/l_j^2$. I used Eqn S22a for stem hydraulic conductance ($K_S$), and Eqn S22b for root hydraulic conductance ($K_R$), treating $k'_S$ and $\kappa_R$ as imposed constants and using values estimated as described below (see *Parameter estimation*). Equation S22b assumes that adding C to the fine root pool results in increased total conducting area, rather than length. Equations S22a and S22b are Eqns A18 and A19 in the Appendix of the main text, respectively. For leaves, I instead used an expression that can be easily constrained by published operational measurements of leaf hydraulic conductance ($K_{\text{leaf}}$, mol $m^{-2}$ $s^{-1}$ $MPa^{-1}$):

(S23) $$K_L = LK_{leaf} \ ,$$

where *L* is leaf area. Equation S23 is Eqn A20 in the Appendix of the main text.

---

***Notes S5. Derivation of expressions for water use in each module***

The transpiration rate of each module is approximately

$$E_{t(m)} \approx \frac{\psi_{soil} - \psi_{L(m)}}{\left(\frac{K_R}{2}\right)^{-1} + K_{S(m)}^{-1} + K_{L(m)}^{-1}} \quad \text{(S24)}$$

where $\psi_{soil}$ is soil water potential and the subscript ($m$) denotes module $m$. The root hydraulic conductance is divided by two to indicate that half of the root water transport capacity is available to each module. More realistically, the entire root transport capacity is shared by both modules, such that the water potential at the point where the two modules diverge, $\psi_D$, is

$$\psi_D = \psi_{soil} - \frac{E_{t1} + E_{t2}}{K_R} \quad \text{(S25)}$$

and the water potential in each module is

$$\psi_{L(m)} = \psi_D - \frac{E_{t(m)}}{K_{(m)}} \quad \text{(S26)}$$

where $K_{(m)} = (K_{S(m)}^{-1} + K_{L(m)}^{-1})^{-1}$. Combining S25 and S26 gives

$$E_{t1} + E_{t2} = K_R(\psi_{soil} - \psi_D) = K_1(\psi_{soil} - \psi_{L1}) + K_2(\psi_{soil} - \psi_{L2}) \quad \text{(S27)}$$

Solving S27 for $\psi_D$ gives

$$\psi_D = \frac{K_R\psi_{soil} + K_1\psi_{L1} + K_2\psi_{L2}}{K_R + K_1 + K_2} \quad \text{(S28)}$$

and applying S28 to S26 (for module #1) gives

(S29) $$E_{t1} = \frac{K_1\big(K_R(\psi_{soil} - \psi_{L1}) + K_2(\psi_{L2} - \psi_{L1})\big)}{K_R + K_1 + K_2}.$$

This is Eqn A17a in the Appendix of the main text. Applying S28 to S26 for module #2 gives

(S30) $$E_{t2} = \frac{K_2\big(K_R(\psi_{soil} - \psi_{L2}) + K_1(\psi_{L1} - \psi_{L2})\big)}{K_R + K_1 + K_2}.$$

which is Eqn A17b in the main text.

***Notes S6. Derivation of expression for leaf temperature***

Leaf energy balance requires that

(S31) $$Q + \mathrm{IR}_{net} - C - \lambda E = 0 \ ,$$

where $Q$ is absorbed shortwave radiation, $\mathrm{IR}_{net}$ is net infrared (longwave) exchange, $C$ is sensible heat loss, and $\lambda E$ is latent heat loss ($\lambda$ is latent heat of vaporization and $E$ is transpiration rate). Convective exchange is $c_{pa} \cdot g_{bh}(T_L - T_A)$, where $c_{pa}$ is heat capacity of the air and $g_{bh}$ is total (2-sided) boundary layer conductance to heat. I assumed that there was no net IR exchange at the lower leaf surface (i.e., that the leaf in question and canopy elements below it had similar temperatures). Outgoing and incoming IR fluxes in the absence of attenuation by other canopy elements are $\varepsilon_L \cdot \sigma_B \cdot T_{LK}^4$ and $\varepsilon_a \cdot \sigma_B \cdot T_{AK}^4$, respectively, where $\varepsilon_L$ and $\varepsilon_a$ are leaf and atmospheric emissivities, respectively, $\sigma_B$ is the Stefan-Boltzmann constant, and $T_{LK}$ and $T_{AK}$ are leaf and air temperatures in kelvins, respectively. I assumed that net IR exchange between the leaf and atmosphere was attenuated by canopy elements above the leaf in question, such that the fraction of net exchange was equal to the fraction of diffuse visible light penetration at an equivalent canopy depth (Leuning *et al.*, 1995), or $\mathrm{IR}_{net} = \mathrm{IR}_{net,top} \cdot \exp(-k_d \cdot \mathrm{LAI})$, where $\mathrm{IR}_{net,top} = \varepsilon_a \cdot \sigma_B \cdot T_{AK}^4 - \varepsilon_L \cdot \sigma_B \cdot T_{LK}^4$, LAI is the cumulative leaf area index above the target leaf, and $k_d$ is the extinction coefficient for diffuse radiation (0.78 for a spherical leaf angle distribution; de Pury & Farquhar, 1997). To estimate LAI for each module, I assumed that overall light attenuation by the canopy above the target leaf occurred with an extinction coefficient $k_i$, such that $i = i_{top} \cdot \exp(-k_i \cdot \mathrm{LAI})$, where $i_{top}$ is the incident visible irradiance above the canopy ($i_{top} = i$ for the sunlit module). Combining these two expressions gives $\mathrm{IR}_{net}/\mathrm{IR}_{net,top} = (i/i_{top})0.78/k_i$. This ratio is termed "$f_{IR}$" in the main text. Thus, $\mathrm{IR}_{net} = f_{IR} \cdot \sigma_B(\varepsilon_a \cdot T_{AK}^4 - \varepsilon_L \cdot T_{LK}^4)$.

Applying these expressions for $\mathrm{IR}_{net}$ and $C$ to Eqn S31 gives

(S32) $$Q + f_{IR}\sigma_B(\epsilon_A T_{AK}^4 - \epsilon_L T_{LK}^4) - c_{pa} g_{bh}(T_L - T_A) - \lambda E = 0 \ .$$

I estimated $T_{LK}$ by expanding $T_{LK}^4 = (T_{AK} + \Delta T)^4$, where $\Delta T = T_{LK} - T_{AK}$), and assuming terms in $\Delta T$ with 2nd and higher powers were negligible:

(S33) $$T_{LK}^4 = (T_{AK} + \Delta T)^4 = T_{AK}^4 + 4T_{AK}^3\Delta T + 6T_{AK}^2\Delta T^2 + 4T_{AK}\Delta T^3 + \Delta T^4 \approx\ T_{AK}^4 + 4T_{AK}^3\Delta T,$$

which is applied to S32 to give

(S34) $$Q + f_{IR}\sigma_B\left(\epsilon_A T_{AK}^4 - \epsilon_L(T_{AK}^4 + 4T_{AK}^3\Delta T)\right) - c_{pa}g_{bh}(T_L - T_A) - \lambda E = 0\ .$$

Solving S34 for $\Delta T$ and noting $T_{LK} = T_{AK} = T_L - T_A$ gives

(S35) $$T_L = T_A + \frac{Q + f_{IR}(\epsilon_A - \epsilon_L)\sigma_B T_{AK}^4 - \lambda E}{4f_{IR}\epsilon_L\sigma_B T_{AK}^3 + c_{pa}g_{bh}}.$$

I assumed incident shortwave radiation (visible + near-infrared [NIR]) was 0.5666·*i*, for radiation in J m$^{-2}$ s$^{-1}$ and incident visible irradiance (photosynthetically active photon flux, PPFD) (*i*) in µmol m$^{-2}$ s$^{-1}$; the value 0.5666 is the ratio of total shortwave energy to photosynthetic photon flux in solar radiation reaching the Earth (de Pury & Farquhar, 1997). Assuming shortwave radiation is half visible and half NIR (Leuning et al. 1995), with leaf absorptances of 0.2 for NIR (Gates *et al.*, 1965) and $\alpha$ for visible light, it follows that $Q = (0.5\alpha + 0.5\cdot0.2)i$. Other factors in Eqn S35 are as follows: $\varepsilon_A = 0.642\cdot(P_A w_A/T_{AK})^{1/7}$ (Leuning *et al.*, 1995), where $P_A$ and $w_A$ are total pressure (101325 Pa) and water vapor mole fraction (mol mol$^{-1}$) of the atmosphere, respectively; $\varepsilon_L = 0.97$; $\sigma_B = 5.67\cdot10^{-8}$ J m$^{-2}$ s$^{-1}$ K$^{-4}$; $\lambda = 4.4\cdot10^4$ J mol$^{-1}$; and $c_{pa} = 29.2$ J mol$^{-1}$ K$^{-1}$.

Note that, although leaf temperature is also influenced by *E* in Eqn S35, the model presented in the main text provides an independent constraint on *E* (from water potentials and hydraulic conductances; Eqns S29 and S30), which can be applied to Eqn S35. This ignores effects of leaf temperature on viscosity, which influences leaf hydraulic conductance; to account for such effects would require iterative numerical solution, increasing computational demand by an order of magnitude. I used air temperature to account for such effects (by multiplying $k'_S$ and $\kappa_L$ by the quantity $(T_{AK}/298.15)^7$, because viscosity increases as the 7th power of temperature). The error incurred by this assumption is probably small, given that predicted differences between leaf and air temperature were predicted to be less than 1K in all simulations (and more importantly, the differences in leaf temperature between the two modules were even smaller).

---
